# Changes in the Anopheles arabiensis transcriptome and gut microbiota profiles associated with Microsporidia MB

**DOI:** 10.1101/2025.06.23.661020

**Authors:** Jacqueline Wahura Waweru, Nicola Mulder, Cynthia Nyambura King’ori, Irene Muiruri, Mwatum Maloba Alakonya, Edward Edmond Makhulu, Lilian Mbaisi Ang’ang’o, Joseph Gichuhi, Daniel Masiga, Jeremy Keith Herren

**Affiliations:** International Centre of Insect Physiology and Ecology (icipe); Computational Biology Division, Department of Integrative Biomedical Sciences, IDM University of Cape Town

## Abstract

*Microsporidia MB* is an endosymbiotic microbe found in *Anopheles* mosquito populations. This symbiont can block *Plasmodium* transmission in mosquitoes, and it can spread through mosquito populations and be sustained over generations by vertical and horizontal transmission. These characteristics make *Microsporidia MB* a potential candidate for symbiont-based malaria vector control. However, the mechanistic basis of interactions between *Microsporidia MB* and *Anopheles arabiensis* is poorly characterized. We investigated how the presence of *Microsporidia MB* affects transcriptomics profiles and the gut microbiota composition and diversity in non-blood-fed and blood-fed (24, 48 and 72 hours post blood meal) *An. arabiensis* mosquitoes. We observed that mosquito infection with *Microsporidia MB* upregulated *farnesoic acid O-methyltransferase,* a gene linked to juvenile hormone biosynthesis in the non-blood-fed mosquitoes. In addition, blood feeding in *Microsporidia MB* positive mosquitoes was associated with an activation of immune-related genes 24 hours after a blood meal, where several genes including the *lipopolysaccharide tumor necrosis factor* gene were upregulated. Interestingly, we also observed that *Microsporidia MB* positivity was associated with a microbiota shift to favor the proliferation of microbes including *Pseudomonas* and *Serratia* 24 hours post blood meal. There were indications of immune system activation in *Microsporidia MB* positive mosquitoes up to 48 hours after a blood meal where factors such as the *peptidoglycan recognition SC2-like* and *lysozyme c-1* were upregulated. Notably, an upregulated immune system at this time point was associated with downregulation of genes associated with metabolism and the restoration of *Serratia*, *Pseudomonas* and other key microbes to relative abundances similar to those recorded in non-blood-fed mosquitoes. At the 72-hour time point, *Microsporidia MB* positive mosquitoes exhibited a downregulation of genes associated with immunity, including *cecropins* and *defensins*, while metabolic processes were predominantly upregulated. Our results provide insights into the effect of *Microsporidia MB* infection on the *An. arabiensis* gene expression and gut microbiota profiles. This work will contribute to mechanistic insights into symbiont-mediated malaria transmission blocking.

## Introduction

Malaria is a primary public health concern, with over 200 million cases and half a million deaths estimated annually. In 2023, 263 million cases in 83 endemic countries were reported, with 94% of the cases being predominant in sub–Saharan Africa. In addition, 597,000 deaths associated with the disease were reported (1). Vector control strategies such as the use of long-lasting insecticidal nets (LLINs) and indoor residual spraying (IRS) have significantly contributed to reducing the number of malaria cases and deaths (2). However, there has been a rising trend in annual cases since 2016, attributed to malaria diagnosis and treatment failures due to parasite evolution, insecticide resistance development, and spread of new invasive malaria vectors including the *Anopheles stephensi* (1). It is, therefore, essential to develop novel and sustainable tools to effectively reduce the malaria burden.

Many insects are associated with heritable endosymbiotic microbes, which have in many instances been reported to influence vector-pathogen interactions (3). An example is the finding that the endosymbiont *Wolbachia* can successfully block the transmission of the arboviral disease Dengue. This finding led to the development of strategies that involve the practical application of symbiotic microbes for blocking vector-borne infectious disease transmission (4–6). Endosymbionts could equally be harnessed to manage other vector-borne infectious diseases like malaria. Notably, the naturally occurring *Microsporidia MB* in *An. arabiensis* mosquitoes, was shown to impair the transmission of *Plasmodium falciparum* (7).

Previous studies have shown that *Microsporidia MB* is avirulent and is vertically and horizontally transmitted through mosquito populations (7,8). These modes of transmission have been hypothesized to aid the ability of the symbiont to spread and be maintained in *Anopheles* mosquito populations (7). In addition, it has exhibited minimal fitness costs on its associated hosts and demonstrated specialized localization patterns within the mosquito, as it is predominantly found in the ovaries and testes and to a lesser extent the midgut (8,9). Further, the endosymbiont is also present in other malaria transmitting vectors other than *An. arabiensis* such as *An. funestus, An. gambiae,* and *An. coluzzii* (8,10). Altogether, many of these characteristics support the possibility of using *Microsporidia MB* as a novel strategy for malaria transmission blocking.

Despite the progress, other facets of the interaction between *Microsporidia MB* and its host, in particular how *Microsporidia MB* impacts host gene expression, remain poorly investigated. By studying these interactions, it may ultimately be possible to elucidate the mechanistic basis of *Microsporidia MB-Anopheles* host interactions and *Microsporidia MB*-*Plasmodium* transmission blocking. In the context of endosymbiont-based transmission blocking, different transmission blocking mechanisms have been proposed, including endosymbionts priming the insect immune system (11), competition between the symbiont and the parasitic agent for important host cellular components (12), and the symbiont encoding factors that are termed as toxins to the parasitic infectious agent (13). It is also possible that endosymbionts can modulate the gut microbiota composition in its insect host in a manner that is less conducive to pathogens. Understanding whether *Microsporidia MB* directly or indirectly impairs the transmission of *Plasmodium* in *An. arabiensis* is essential for predicting the long-term success of *Microsporidia MB-*based malaria disease control strategy. In this study, we hypothesized that *Microsporidia MB* impairs *Plasmodium* development by (I) activating mosquito immune responses, and (II) restructuring the gut microbiota to favor anti-*Plasmodium* bacteria.

In *Anopheles* mosquitoes, blood feeding is known to trigger changes in the gut physiology, immunity, and microbiome (14,15). These changes, particularly within the first 72 hours post-blood meal, play a critical role in determining the ability of *Plasmodium* parasites to invade and establish in the mosquito midgut (16). Therefore, this study investigated host gene expression and gut microbiota dynamics in *Microsporidia MB*-positive and -negative *Anopheles* mosquitoes at key time points: 24-, 48-, and 72-hours post-blood feeding, as well as in seven-day-old non-blood-fed individuals. To control for potential genetic background effects, *Microsporidia MB*-negative pooled siblings of *Microsporidia MB*-positive mosquitoes were also included.

Previously, it was shown that the rate of *Microsporidia MB* vertical transmission ranges between 45-100% (average of 62%), indicating that not all offspring from a *Microsporidia MB* infected G_0_ *An. arabiensis* females end up positive for the endosymbiont as adults, hence it is possible to have negative siblings from positive G_0_ *An. arabiensis* females (7,17). Recent evidence suggests that *Microsporidia MB* negative siblings are likely to have cleared *Microsporidia MB* infections, since almost all the eggs laid by *Microsporidia MB* infected G_0_ *An. arabiensis* females are known to be infected (7). It is possible that clearing *Microsporidia MB* leads to changes in gene expression. In addition, rearing *Microsporidia MB* positive and negative sibling mosquitoes in the same larval trays likely results in similar microbiota profiles. We therefore investigated the differences in gene expression and microbiota profile between *Microsporidia MB* positive and negative pooled siblings as well as between *Microsporidia MB* positive and negative pooled non-sibling mosquitoes, reared in different trays. To better understand interactions between *Microsporidia MB* at the tissue level we profiled changes in gene expression in two different tissues, the gut, and the fat body. Our findings highlight a variety of differences between *Microsporidia MB* infected and uninfected mosquitoes which shed light on the nature of the symbiotic relationship and the possible basis of the transmission blocking phenotype.

## RESULTS

### Principal component analysis revealed different gene clustering patterns between *Microsporidia MB*-positive and -negative pooled sibling and non-sibling mosquitoes

Cumulatively, 34,804,584 reads were recovered from the *RNAseq* data analysis. This data was analysed per time point and tissue to address time and tissue dependent expression changes. Principal component analysis (PCA) of all genes from *Microsporidia MB* positive, -negative pooled sibling and non-sibling mosquitoes (Figure 1A), and constituting the non-blood-fed and blood-fed mosquitoes (24, 48, and 72 hours post blood meal) indicated that genes from the guts of *Microsporidia MB*-positive mosquitoes clustered more closely with those from *Microsporidia MB*-negative pooled sibling mosquitoes, and separately from *Microsporidia MB*-negative pooled non-sibling mosquitoes (Figure 1B, C, D and E). The close gene clustering patterns between *Microsporidia MB*-positive and -negative pooled sibling mosquitoes resulted in fewer differentially expressed genes (DEGs) (Supplementary Table S1, S2, S3 and S4), compared to the numbers recorded when *Microsporidia MB*-positive mosquitoes were compared to *Microsporidia MB*-negative pooled non-sibling mosquitoes (Supplementary Table S5, S6, S7 and S8).

**Figure 1:**
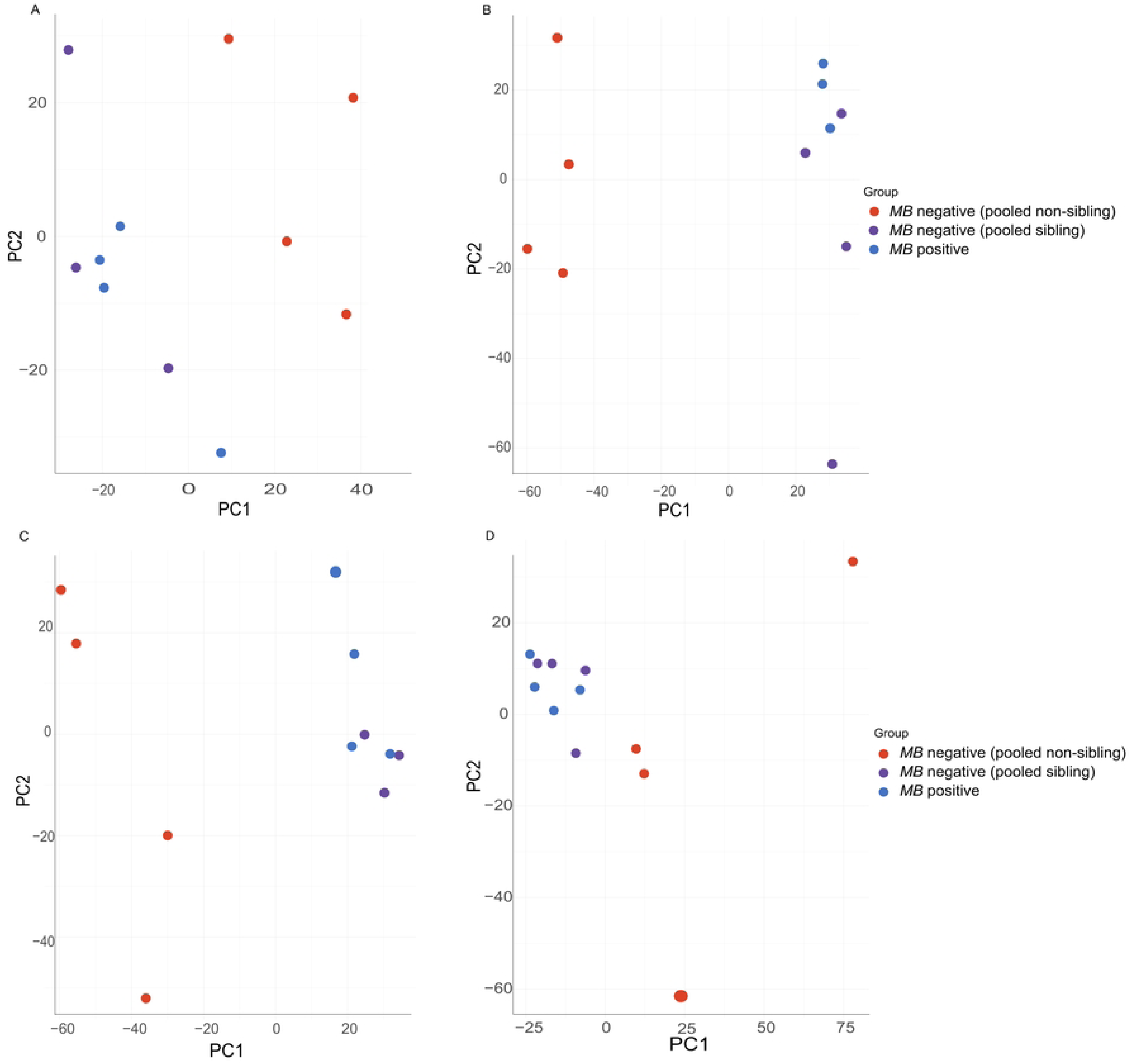
(A) A diagram representing different categories of *An. arabiensis* mosquitoes used in our study. B, C, D and E are representative PCAs to demonstrate the clustering patterns of genes in the guts of *Microsporidia MB* positive and -negative pooled sibling and pooled non-sibling mosquito guts across different time points. (B) represents the clustering patterns in non-blood fed mosquitoes, (C) 24 hours post blood meal, (D) 48 hours post blood meal and (E) 72 hours post a blood meal. Blue represents *Microsporidia MB* positive mosquitoes, purple -negative sibling mosquitoes and red -negative non-sibling mosquitoes. Red represents upregulation while blue represents downregulation.

Similarly, genes from the fat bodies from *Microsporidia MB*-positive mosquitoes in both the blood-fed and non-blood-fed categories clustered more closely with those from *Microsporidia MB*-negative pooled sibling mosquitoes and separately from the *Microsporidia MB*-negative pooled non-sibling mosquitoes (Supplementary Figure S1A, B, C, D). In addition, a comparison between *Microsporidia MB*-positive and the *Microsporidia MB*-negative pooled sibling mosquitoes yielded fewer DEGs (Supplementary Table S9, S10, S11, S12), compared to the numbers observed when *Microsporidia MB*-positive mosquitoes were compared to *Microsporidia MB*-negative pooled non-sibling mosquitoes (Supplementary Table S13, S14, S15, S16). The few DEG counts when *Microsporidia MB*-positive mosquitoes were compared to the -negative pooled sibling mosquitoes led to their gene ontology enrichment analysis in both the guts and fat body not being possible.

### Differentially expressed genes associated with *Microsporidia MB*-positive mosquitoes compared to *Microsporidia MB*-negative sibling and non-sibling mosquitoes across time points

#### Non-blood fed mosquitoes

We identified 36 DEGs when the guts from non-blood-fed *Microsporidia MB*-positive and -negative pooled sibling mosquitoes were compared, 23 of which were upregulated and 13 downregulated in *Microsporidia MB*-positive mosquitoes (Supplementary table S1). Upregulated genes included *farnesoic acid O-methyltransferase*, *glutathione S transferase 1-1 like*, and *chymotrypsin 2 like* while *trypsin 4,6 and 7*, *chymotrypsin B1* and *farnesol dehydrogenase* were downregulated (Supplementary table S1). When guts from *Microsporidia MB*-positive mosquitoes were compared to those from *Microsporidia MB*-negative pooled non-sibling mosquitoes, 34 genes were differentially expressed, with *chymotrypsin 2 like*, *serine protease SP24D like*, *farnesoic acid O-methyltransferase*, *farnesol dehydrogenase* and *glutathione S transferase* being upregulated and *trypsin 4,6 and 7*, and *chymotrypsin B1* like being downregulated in *Microsporidia MB*-positive mosquitoes (Figure 2A, Supplementary table S5).

**Figure 2:**
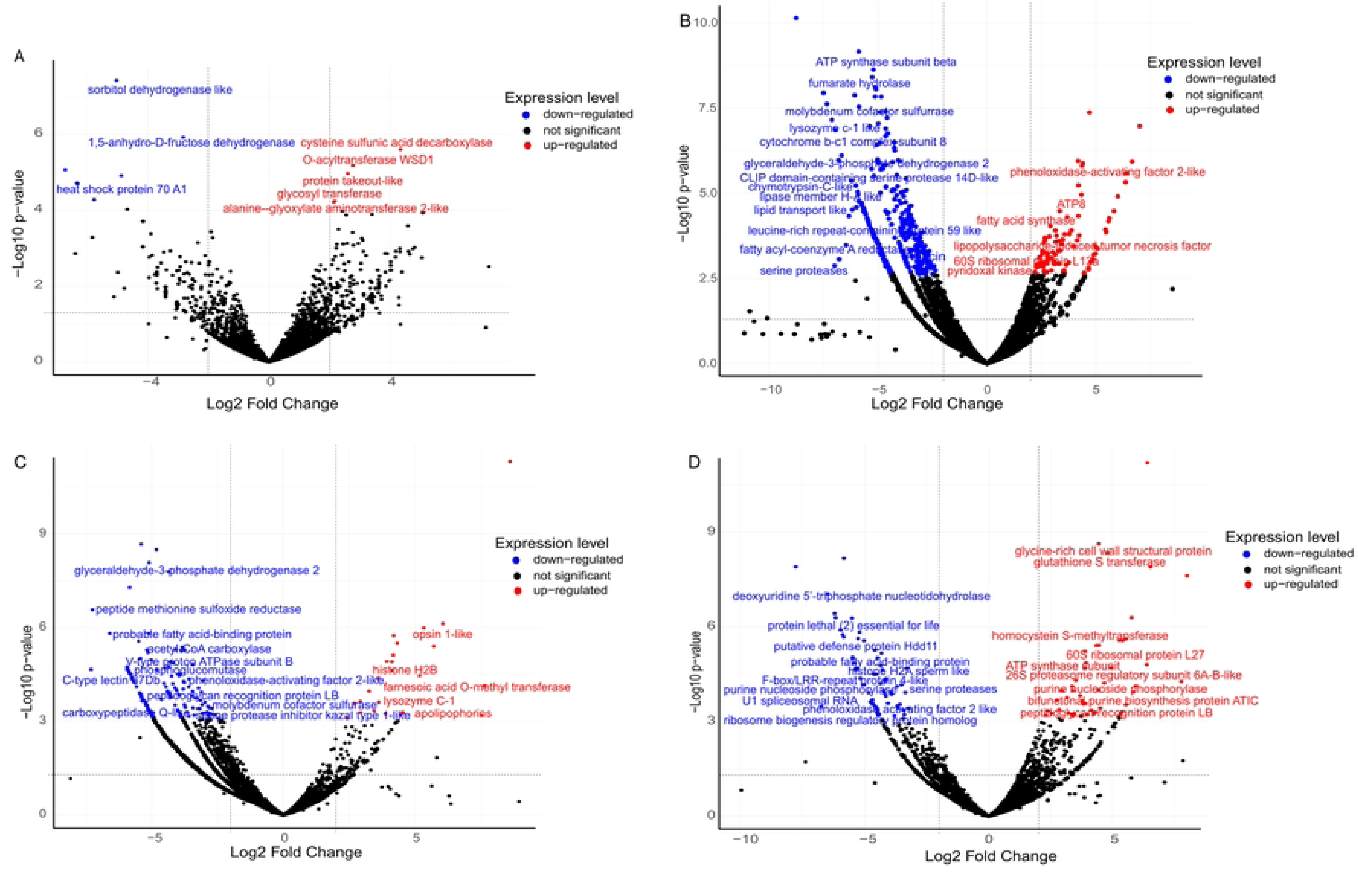
Volcano plots showing the differentially expressed genes in the guts of *Microsporidia MB* positive mosquitoes compared to -negative pooled non-sibling mosquitoes. A, B, C and D represent the differentially expressed genes in non-blood-fed mosquitoes, 24 hours post blood meal, 48 hours post blood meal and 72 hours post blood meal respectively. Red represents upregulation while blue represents downregulation.

In the fat body, a comparison of *Microsporidia MB*-positive and -negative pooled sibling mosquitoes yielded only one significant and upregulated gene, *cysteine sulfunic acid decarboxylase* (Supplementary table S9). In contrast, a comparison of *Microsporidia MB*-positive and -negative pooled non-sibling mosquitoes yielded 14 DEGs, 5 of which were upregulated and 9 downregulated in *Microsporidia MB*-positive mosquitoes (Supplementary table S13). Upregulated genes included *glycosyl transferase*, *cysteine sulfunic acid decarboxylase* and *alanine glyoxylate aminotransferase* while *heat shock protein*, *sorbitol dehydrogenase* and *1,5-anhydro-D-fructose reductase* were two of the downregulated genes (Supplementary figure S2A, Supplementary table S13).

#### 24 hours post blood meal

Ten genes were differentially expressed, 3 of which were upregulated and 7 downregulated when guts from *Microsporidia MB*-positive and -negative pooled sibling mosquitoes were compared 24 hours post blood meal (Supplementary table S2). The downregulated genes included *glycine-rich cell wall structural protein*, *cell wall integrity* and *stress response component 4-like*, and *general transcriptional corepressor trfA-like* while upregulated genes included *urease accessory protein* and *peroxisomal multifunctional enzyme type 2 like* (Supplementary table S2). A comparison of guts from *Microsporidia MB*-positive and -negative pooled non-sibling mosquitoes on the other hand yielded 145 differentially expressed genes, 31 of which were upregulated and 114 downregulated in *Microsporidia MB*-positive mosquitoes at this time point (Supplementary table S6). Upregulated genes included the *peptidoglycan recognition protein SC*, *lipopolysaccharide tumor necrosis factor (LITAF)*, *lysozyme C-1*, *urease accessory protein* and *oxidative stress-related genes*, including *glutaredox C4* and *peroxiredoxin 6-like,* while downregulated genes included the *molybdenum cofactor sulfurase*, *serine palmitoyl transferase*, and *alanine glyoxylate aminotransferase* (Figure 2B, Supplementary table S6). Interestingly, only *urease accessory protein* was upregulated when guts from *Microsporidia MB*-positive were compared to those from both sets of controls at this time (Supplementary table S2 and S6).

When the fat bodies from *Microsporidia MB*-positive mosquitoes were compared to *Microsporidia MB-*negative pooled sibling mosquitoes, 42 genes were differentially expressed, with 2 being upregulated and 40 downregulated in *Microsporidia MB*-positive mosquitoes (Supplementary table S10). *Chymotrypsin inhibitor like* was upregulated while *glutathione S transferase*, *flightin*, *phosphate carrier protein*, *acylphosphatase*, *vesicle transport protein*, *ADP, ATP carrier protein* and *myofilin* were downregulated (Supplementary table S10). Further, we identified 448 DEGS when fat bodies from *Microsporidia MB*-positive mosquitoes were compared to those from *Microsporidia MB*-negative pooled non-sibling mosquitoes (Supplementary table S14). Intriguingly, *LITAF*, alongside *phenoloxidase activating factor 2 like*, which were upregulated in the guts when *Microsporidia MB*-positive mosquitoes were compared to *Microsporidia MB-*negative pooled non-sibling mosquitoes, were also upregulated in the fat bodies when *Microsporidia MB*-positive and -negative pooled non-sibling mosquitoes were compared (Supplementary figure S2B and Supplementary table S14). Interestingly, *glutathione S transferase*, *flightin, phosphate carrier protein*, *acylphosphatase*, *vesicle transport protein*, *ADP, ATP carrier protein* and *myofilin* were also all downregulated in the fat bodies of *Microsporidia MB-*positive mosquitoes compared to -negative pooled non-sibling mosquitoes (Supplementary table S14).

#### 48 hours post blood meal

No significant genes were identified at this time point when guts from *Microsporidia MB*-positive mosquitoes were compared to those from *Microsporidia MB*-negative pooled sibling mosquitoes (Supplementary table S3). Interestingly, 537 genes were differentially expressed in the guts, 50 of which were upregulated, and 488 downregulated when *Microsporidia MB*-positive mosquitoes were compared to *Microsporidia MB*-negative pooled non-sibling mosquitoes (Supplementary Table S7). The *peptidoglycan recognition SC2-like* and *lysozyme c-1*, remained upregulated even at this time point while *leucine-rich melanocyte differentiation associated protein-like*, and *galectin 4* were downregulated. Metabolism-related genes such as *glucose dehydrogenase*, *alpha aminoapidic semialdehyde synthase*, *glyceraldehyde-3-phosphate dehydrogenase 2*, and *probable NADH dehydrogenase* among others, were generally downregulated (Figure 2C, Supplementary Table S7).

Fat bodies from *Microsporidia MB*-positive compared to those from *Microsporidia MB-* negative pooled sibling mosquitoes, only yielded 5 differentially expressed genes, with only 1 upregulated and 4 downregulated (Supplementary table S11). The downregulated included *cecropin-B*, *phosphate carrier protein* and *30 kDa salivary gland allergen Aed a 3-like* (Supplementary table S11). 161 genes on the other hand were differentially expressed, with 26 upregulated and 135 downregulated when fat bodies from *Microsporidia MB*-positive were compared to those from *Microsporidia MB-*negative pooled non-sibling mosquitoes (Supplementary table S11). Interestingly, the gene *lysozyme c-1* upregulated in the gut was also upregulated in the fat body when *Microsporidia MB*-positive mosquitoes were compared to *Microsporidia MB-*negative pooled non-sibling mosquitoes, at this time point (Supplementary Figure S2C, Supplementary Table S15). The *phosphate carrier protein* was downregulated in the fat body when a comparison between *Microsporidia MB*-positive and both control types was done (Supplementary table S11 and S15).

#### 72 hours post blood meal

Sixty-one genes were differentially expressed, 57 of which were upregulated and 4 downregulated when guts from *Microsporidia MB*-positive were compared to those from *Microsporidia MB-*negative pooled sibling mosquitoes (Supplementary table S8). Upregulated genes in *Microsporidia MB* infected mosquitoes included *acetyl-CoA carboxylase*, *homocysteine S-methyltransferase*, *molybdenum cofactor sulfurase*, and *fatty acid synthase* among others (Supplementary table S4). A total of 333 genes on the other hand were differentially expressed, with 90 upregulated and 243 downregulated in *Microsporidia MB*-positive mosquitoes compared to *Microsporidia MB-*negative pooled non-sibling mosquitoes (Supplementary table S8). Specifically, genes associated with immunity were downregulated including *defensin*, *cecropin A*, *cecropin B*, *cecropin C*, *LITAF*, *peptidoglycan recognition protein*, and *lysozyme c like* (Figure 2D, Supplementary table S8).

In the fat body, only 2 genes, *ornithine decarboxylase* and *galectin*, were significantly expressed and downregulated when *Microsporidia MB*-positive mosquitoes were compared to *Microsporidia MB-*negative pooled sibling mosquitoes at this time point (Supplementary table S12). Conversely, a comparison of fat bodies from *Microsporidia MB*-positive and -negative pooled non-sibling mosquitoes yielded 117 differentially expressed genes, with *ATP synthase subunit*, *purine nucleoside phosphorylase*, *bifunctional purine biosynthetic protein*, *phosphoenolpyruvate carboxykinase*, being upregulated while *glutathione S transferase*, *phenoloxidase activating factor 2,* among others, were downregulated in *Microsporidia MB*-positive mosquitoes (Supplementary figure S2D, Supplementary Table S16).

### Differentially enriched gene ontology (GO) pathways in *Microsporidia MB*-positive mosquitoes compared to *Microsporidia MB*-negative pooled non-sibling mosquitoes

#### Non-blood-fed mosquitoes

DEGs from non-blood-fed guts were annotated to a range of biological and molecular GO terms, as shown in (Figure 3A). The upregulated pathways included proteolysis, peptidase, inorganic diphosphate activity among others. Downregulated pathways, on the other hand, included protein metabolic, serine endopeptidase, and catalytic activity acting on a protein (Figure 3A, Supplementary table S17). In the fat body, *Microsporidia MB* infection induced upregulation of pathways related to carbohydrate metabolism, vitamin binding, and Vitamin B6 metabolism. In contrast, pathways spanning ribonucleotide binding, pyrophosphatase, and purine ribonucleotide binding remained downregulated in the fat bodies (Supplementary figure S3A, Supplementary table S18).

**Figure 3:**
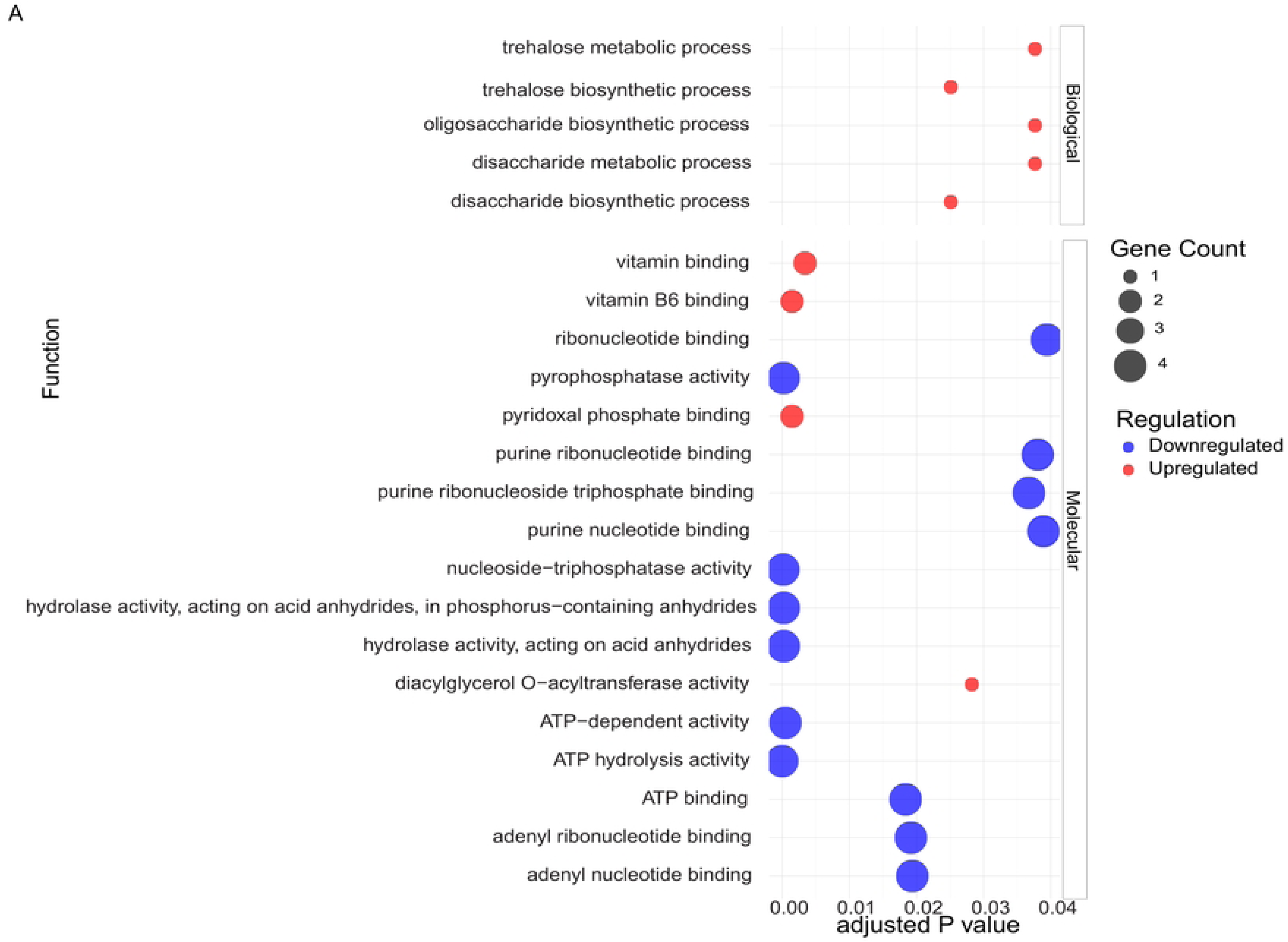
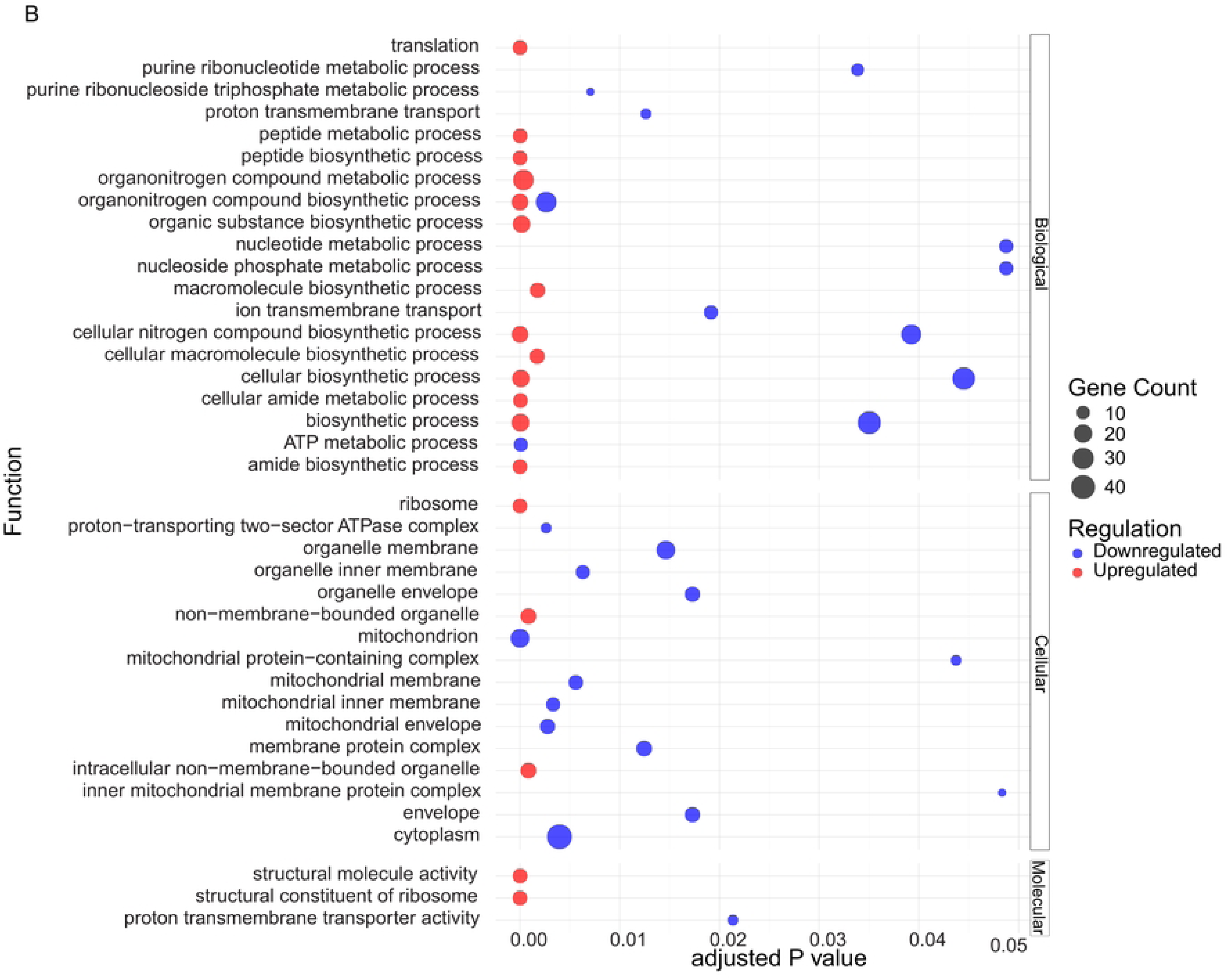
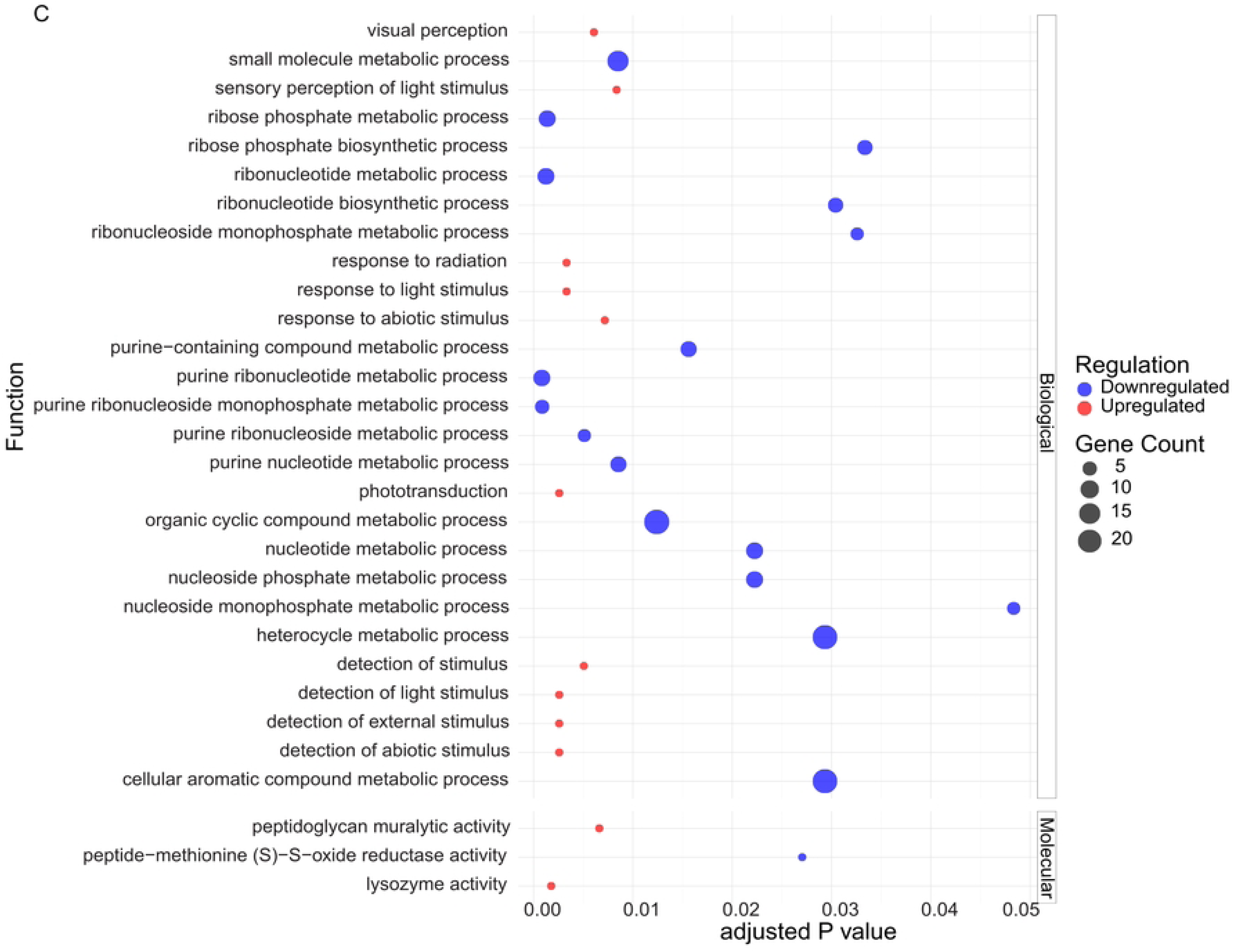
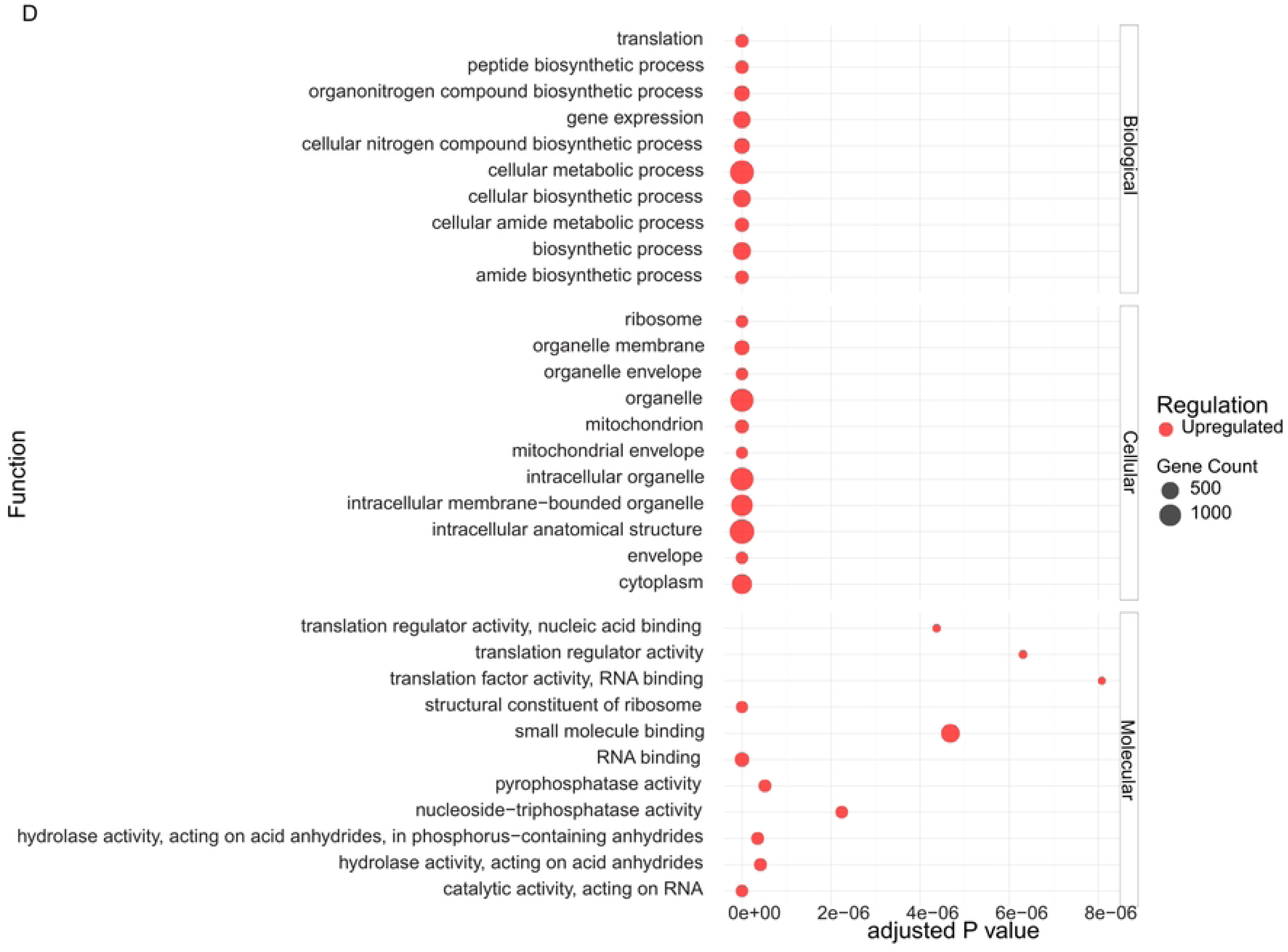
Gene ontology analysis of differentially expressed genes in the guts of *Microsporidia MB* positive compared to -negative pooled non-sibling mosquitoes. A, B, C and D represent the enriched pathways in non-blood-fed mosquitoes, 24 hours post blood meal, 48 hours post blood meal and 72 hours post blood meal respectively. Red represents upregulation while blue represents downregulation.

#### 24 hours post blood meal

Different biological, cellular and molecular GO terms were significant in the guts of *Microsporidia MB* positive mosquitoes at this time point (Figure 3B). The upregulated terms included the peptidoglycan muralytic pathway, while terms associated with metabolism were mostly downregulated, including ribose phosphate metabolic pathway, ribonucleotide metabolic process, organonitrogen compound biosynthetic pathway, carbohydrate derivative pathway, and Vitamin B6 binding pathway (Figure 3B, Supplementary table S19). In contrast to downregulated metabolism in the guts of infected mosquitoes 24 hours post-blood meal, fat bodies recorded an upregulation of metabolic related processes primarily involving amino acid metabolism (Supplementary figure S3B, Supplementary table S20).

#### 48 hours post blood meal

Several biological, cellular, and molecular terms were differentially enriched in the guts at this time point, with the peptidoglycan muralytic activity and chitin-binding pathways being upregulated and most metabolism-related pathways including nucleotide metabolism, organonitrogen compound biosynthetic process, purine ribonucleotide metabolic process, and small molecule metabolic process being downregulated (Figure 3C, Supplementary table S21). In congruence with the observations made in the gut, the fat body also exhibited an upregulation of the peptidoglycan muralytic activity, lysozyme activity, along with downregulation of metabolic pathways including purine ribonucleotide metabolism, small molecule metabolic process, ribose phosphate metabolic process, and ribonucleotide biosynthetic process (Supplementary figure S3C, Supplementary table S22).

#### 72 hours post blood meal

Immune-related GO terms such as humoral immune response and serine-type peptidase activity were downregulated at this time point in the gut (Figure 3D, Supplementary table S23). Interestingly, metabolism related pathways were upregulated in the fat body of *Microsporidia MB* positive mosquitoes, including peptide biosynthetic process, organonitrogen compound biosynthetic process, and cellular amide biosynthetic process (Supplementary Figure S3D, Supplementary table S24).

### Gut bacterial community composition in *Microsporidia MB*-positive mosquitoes

A total of 20,173,248 paired-end reads spanning the V3-V4 region of the bacterial 16s rRNA were sequenced, with per-sample read counts ranging from 104,112 to 663,016. After quality assessment and denoising the data, 7,230,708 reads remained for downstream analysis, yielding 3316 amplicon sequence variants (ASVs). Further filtering to remove unwanted ASVs, including Mitochondria, Chloroplast, Archaea, Cyanobacteria, Chloroflexi, Eukaryota, and those classified as unassigned reads at the kingdom level as well as ASVs with abundances below five, resulted in 2,255 ASVs that were retained for downstream analysis.

At the genus level, 19 genera with an overall abundance higher than 0.5% were classified as representative gut microbiota communities in the mosquitoes being studied (Figure 4A, B and C, Supplementary Table S25). *Rhodococcus* in the phylum Actinobacteria, and *Pseudomonas*, *Allorhizobium*, *Serratia*, and *Achromobacter,* all in the phylum Proteobacteria, were the top 5 most abundant genera with overall relative abundances of 19.23%, 14.71%, 7.92%, 7.31%, and 5.43% respectively. Other highly abundant genera in the dataset included *Brevundimonas, Ralstonia, Acidovorax, Variovorax, Phyllobacterium, Delftia, Mesorhizobium, Aeromonas, Asaia, Acinetobacter,* and *Rahnella* all in the phylum Proteobacteria, *Chryseobacterium* and *Enterococcus* in the phylum Bacteroidota and *Elizabethkingia* in the phylum Firmicutes (Figure 4A, B and C, Supplementary Table S25). Among the most abundant genera, *Rhodococcus*, *Pseudomonas*, *Allorhizobium,* and *Achromobacter* were detected in 100% of the samples. *Acinetobacter, Brevundimonas, Elizabethkingia, Mesorhizobium, Acidovorax, and Delftia* were also highly prevalent across samples, with their prevalence estimates spanning between 90% and 97% (Supplementary Table S26).

**Figure 4:**
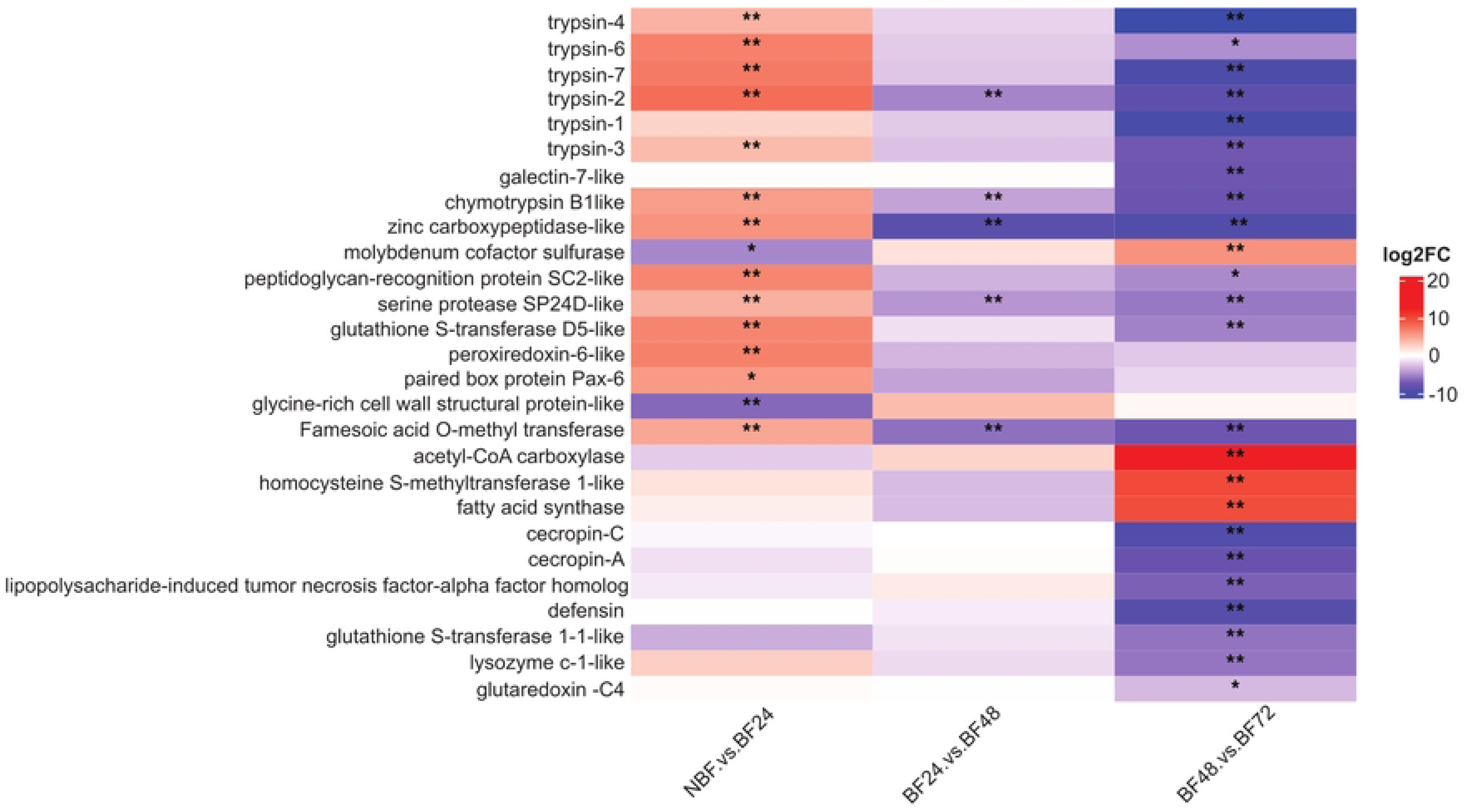
Bacterial genera members in *Microsporidia MB* positive and -negative (-negative pooled sibling mosquitoes in purple and -negative pooled non-sibling mosquitoes in red) *An. arabiensis* mosquitoes. Three time points including (A) 7 days old non-blood-fed and (B) 24 and (C) 48 hours post blood meal among mosquitoes fed on blood 3 days post eclosion were included.

### Effects of *Microsporidia MB* infection on bacterial profiles and densities

Genus-level microbiota composition varied significantly with *Microsporidia MB* infection, especially following blood feeding. *Rhodococcus, Elizabethkingia, Chryseobacterium, Brevundimonas,* and *Mesorhizobium,* for example, had the highest relative abundances in non-blood-fed *Microsporidia MB*-positive compared to *Microsporidia MB*-negative pooled sibling and pooled non-sibling mosquitoes (Supplementary Table S25). However, 24 hours after blood feeding, their relative abundances dropped to the lowest levels in *Microsporidia MB*-positive mosquitoes compared to *Microsporidia MB-*negative pooled sibling and pooled non-sibling mosquitoes, only slightly increasing again 48 hours after blood meal. *Serratia* and *Pseudomonas,* on the other hand, had the lowest relative abundances in non-blood fed *Microsporidia MB*-positive mosquitoes compared to *Microsporidia MB-*negative pooled sibling and pooled non-sibling mosquitoes. The relative abundances of the two later proliferated with blood feeding, attaining the highest relative abundances 24 hours post-blood feeding in the *Microsporidia MB*-positive mosquitoes compared to the *Microsporidia MB-* negative pooled sibling and pooled non-sibling mosquitoes. *Serratia* and *Pseudomonas* dropped 48 hours post-blood meal in *Microsporidia MB*-positive mosquitoes to relative abundances similar to what was observed in non-blood fed *Microsporidia MB*-positive mosquitoes. On the contrary, the *Microsporidia MB-*negative pooled sibling and pooled non-sibling mosquitoes recorded an increased relative abundance in *Pseudomonas* and *Serratia* respectively at this time point (Supplementary Table S25).

An evaluation of the effect of *Microsporidia MB* infection on the gut bacterial density on the other hand showed that *Microsporidia MB* infection resulted in higher bacterial densities in non-blood-fed *Microsporidia MB*-positive mosquitoes compared to the *Microsporidia MB*-negative pooled sibling and pooled non-sibling mosquitoes (Figure 5).

**Figure 5:**
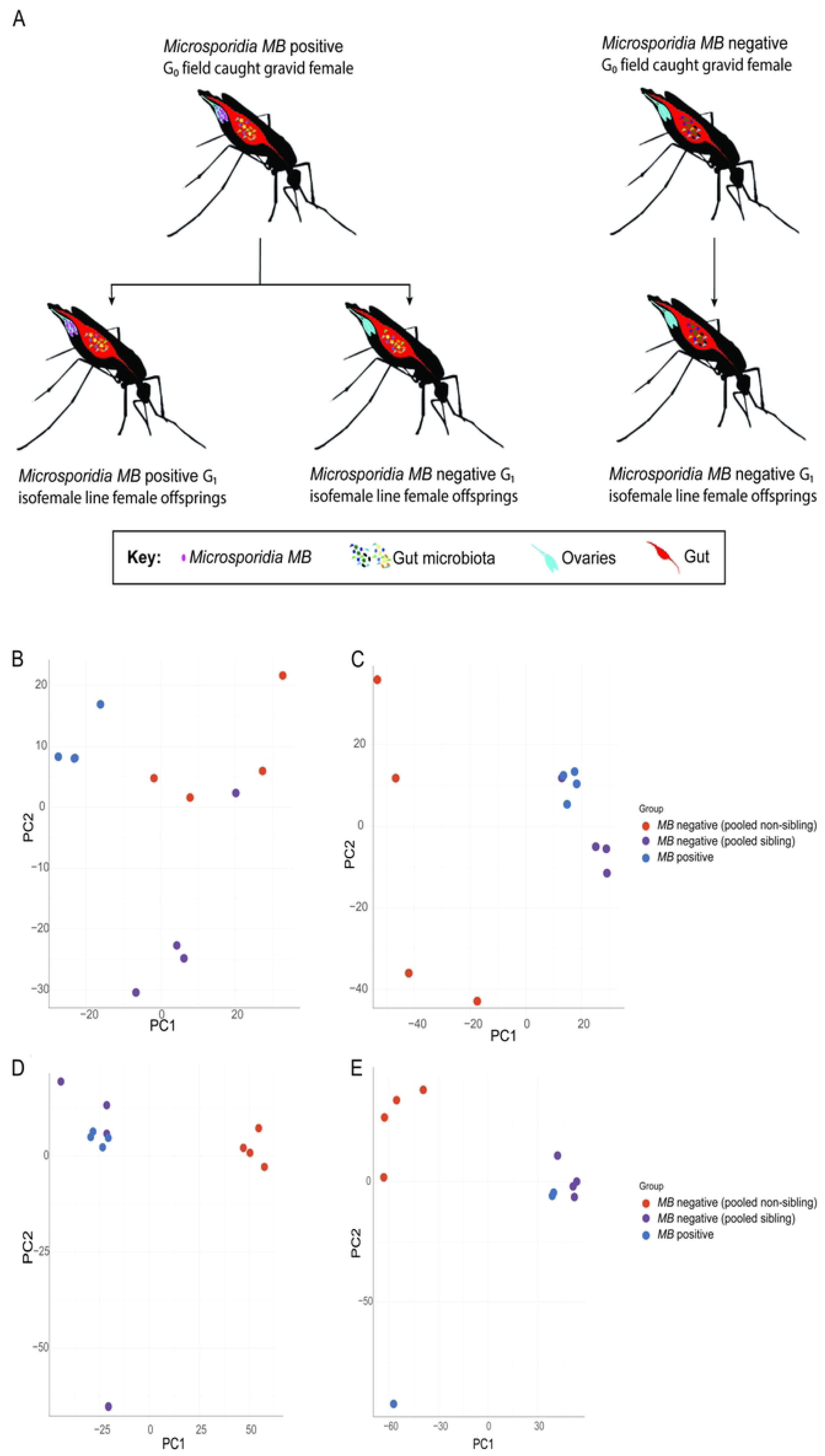
Bacterial densities in non-blood-fed mosquitoes positive and -negative for *Microsporidia MB. Microsporidia MB* positive mosquitoes are in blue, -negative pooled sibling mosquitoes in purple and -negative pooled non-sibling mosquitoes in red.

### Effect of *Microsporidia MB* infection and blood feeding on bacterial diversity

Next, we determined whether mosquito infection with *Microsporidia MB* affected its host’s gut microbiota diversity. Principal coordinate analysis (PCoA) showed that microbiota from *Microsporidia MB*-positive mosquitoes clustered more closely with those from *Microsporidia MB-*negative pooled sibling mosquitoes, and at all time points i.e., non-blood-fed, 24 hours, and 48 hours post blood meal (Figure 6A, B, C, D, E and F). Conversely, microbiota from *Microsporidia MB*-positive mosquitoes clustered distinctly from that of *Microsporidia MB-* negative pooled non-sibling mosquitoes across the various time points. Close clustering between *Microsporidia MB*-positive and -negative pooled sibling mosquitoes and distinct clustering between *Microsporidia MB*-positive and -negative pooled non-sibling mosquitoes were further supported by PERMANOVA pairwise analysis (Table 1, Supplementary Table S27).

**Figure 6.**
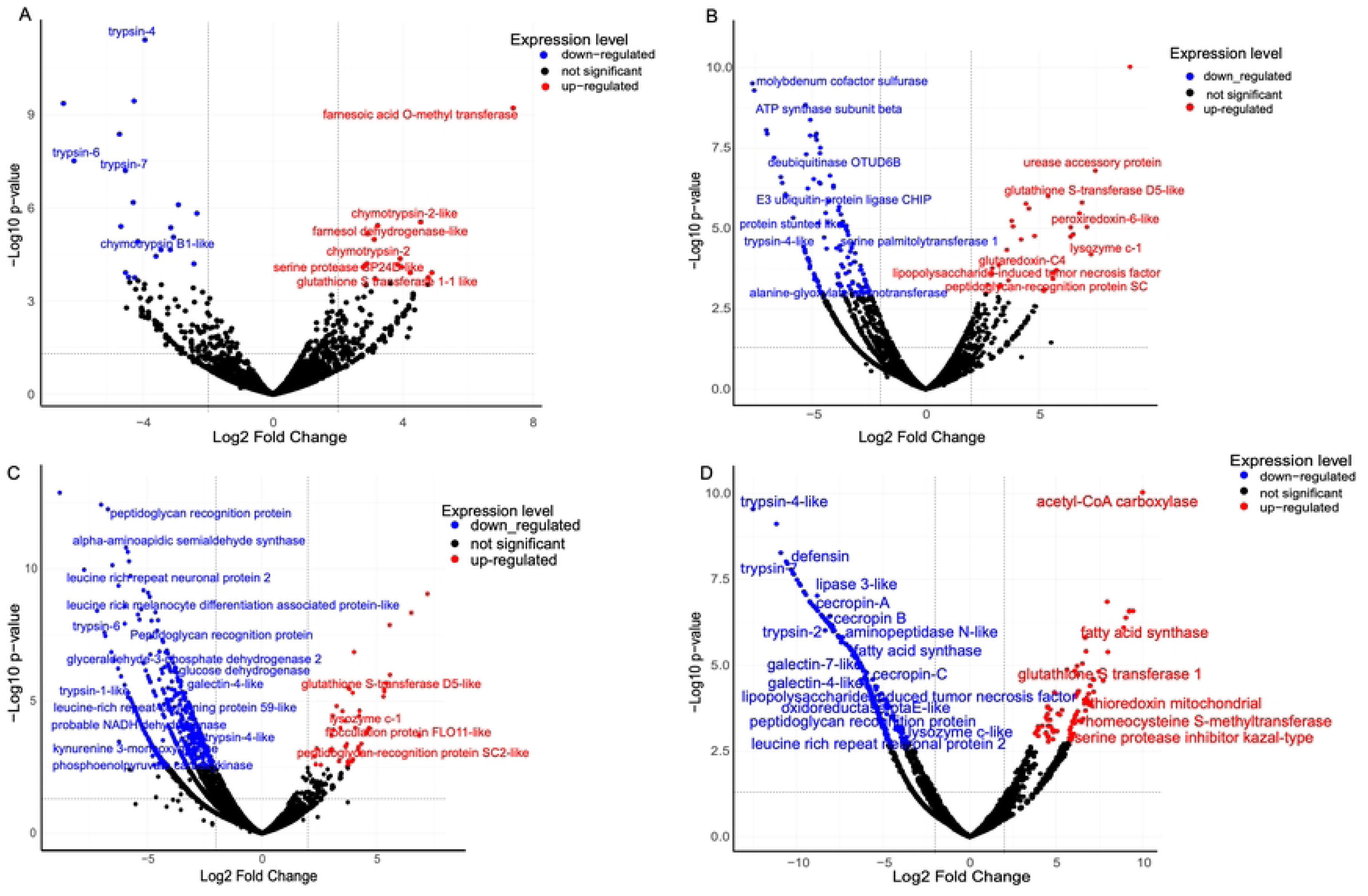
Beta diversity estimates of the gut microbiota data using unweighted unifrac (A, C and E), and non-metric multi-dimensional scaling (B, D and F) from non-blood-fed (A and B) and blood fed (24 (C and D) and 48 (E and F) hours post blood meal) mosquito guts. *Microsporidia MB* positive mosquitoes are represented in blue, negative pooled sibling mosquitoes in purple and negative pooled non-sibling mosquitoes in red.

**Table 1:**
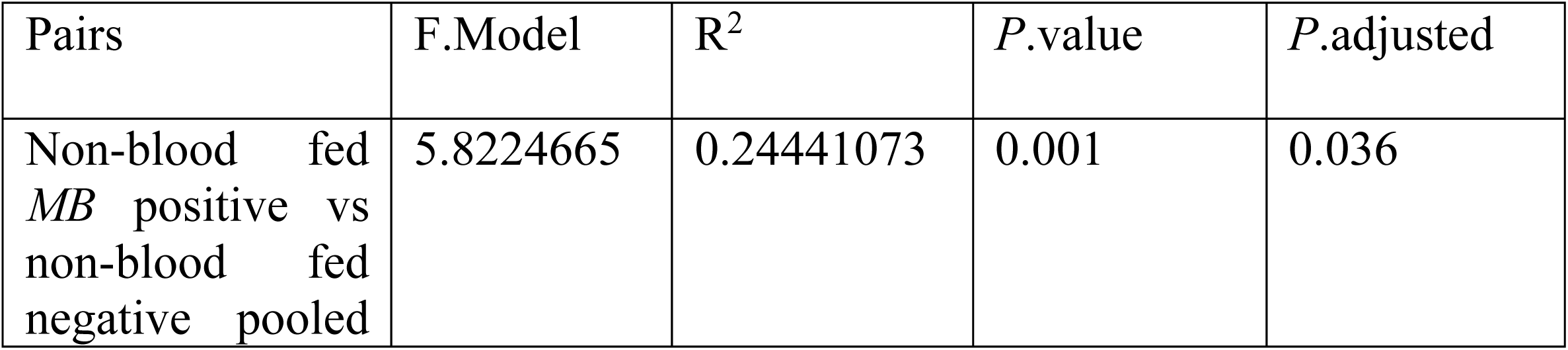
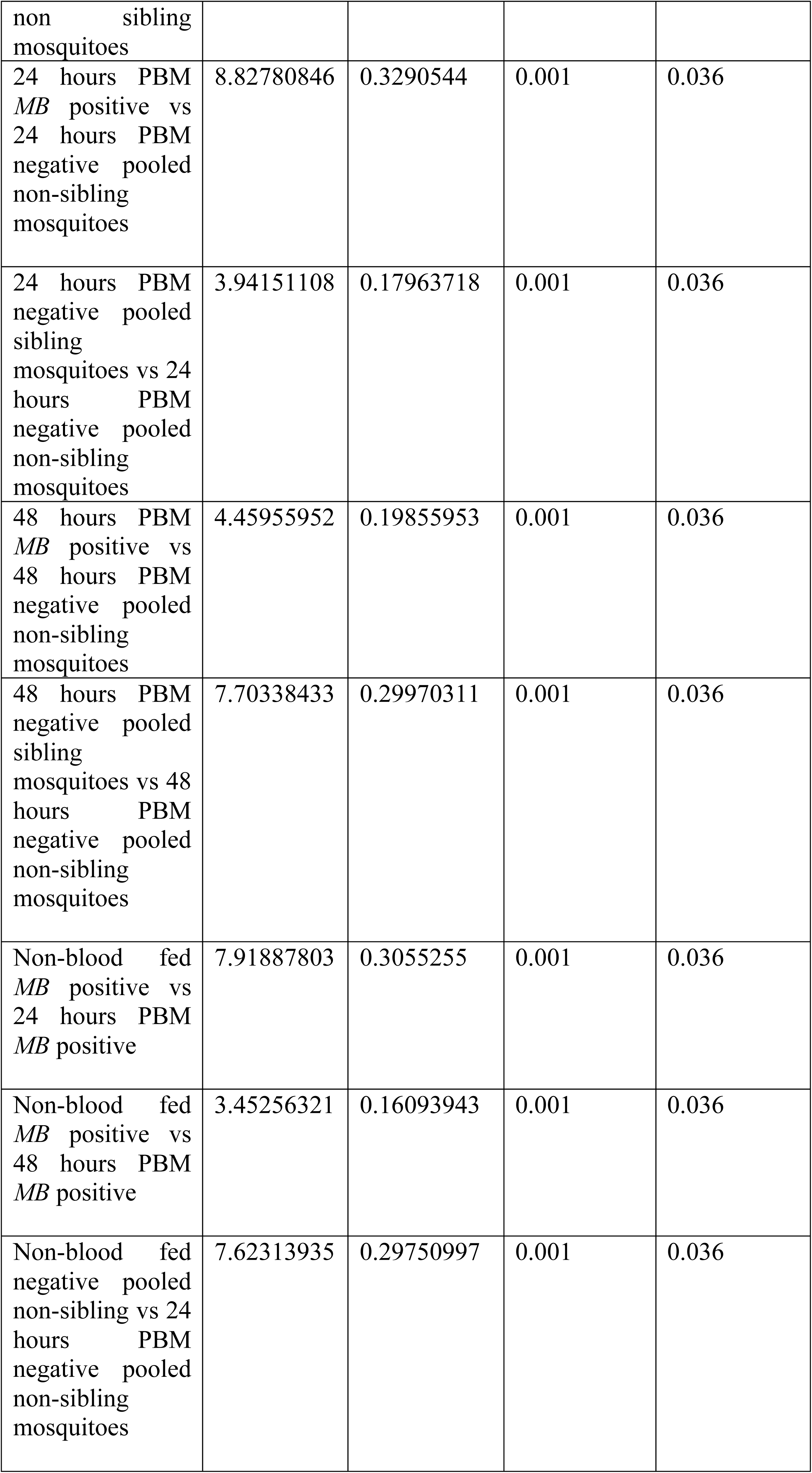
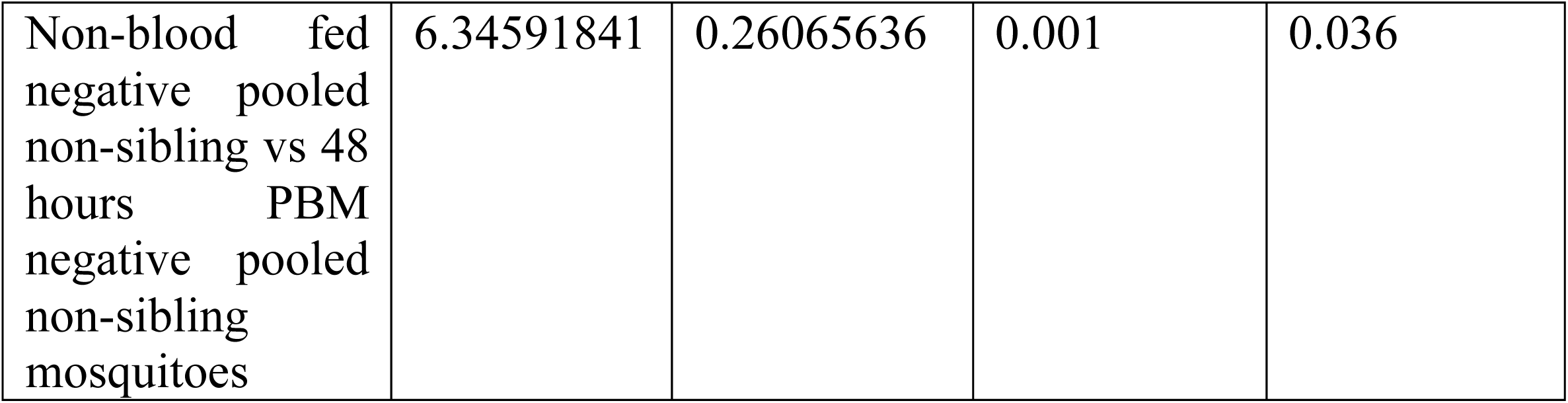
A tabulation of differential clustering statistics as determined using PERMANOVA pairwise analysis.

Further, we observed that microbiota of non-blood-fed *Microsporidia MB*-positive mosquitoes clustered separately from the microbes in blood-fed mosquitoes. Notably, microbes from mosquitoes fed on blood at the 24- and 48-hour time points clustered together, indicating a similar gut microbiota community structure after blood feeding (Figure 7A and B). On the contrary, the negative pooled sibling mosquitoes demonstrated a different clustering pattern from the *Microsporidia MB*-positive and -negative mosquitoes, since both the non-blood-fed and blood-fed mosquitoes clustered together, indicating that blood-feeding did not influence the gut microbiota structure of these mosquitoes (Figure 7C and D). Similarly to *Microsporidia MB*-positive mosquitoes, the *Microsporidia MB*-negative mosquitoes also exhibited a similar clustering pattern, indicating that blood-feeding also influenced the gut microbiota structure of these mosquitoes (Figure 7E and F).The findings that blood-feeding influenced the gut microbiota structure in *Microsporidia MB*-positive and the -negative non-sibling mosquitoes were further supported by PERMANOVA pairwise analysis (Table 1, Supplementary Table S27).

**Figure 7:**
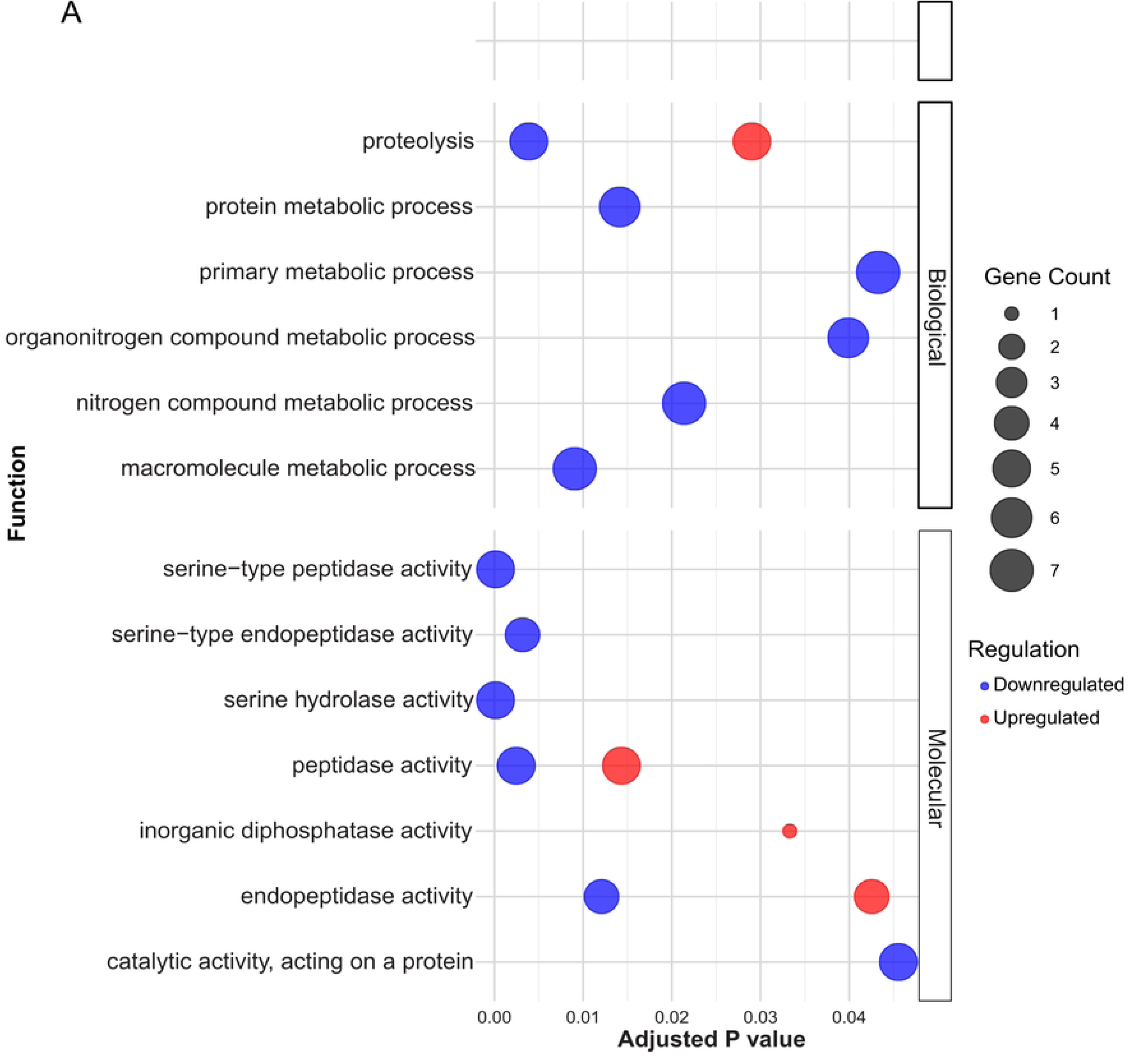
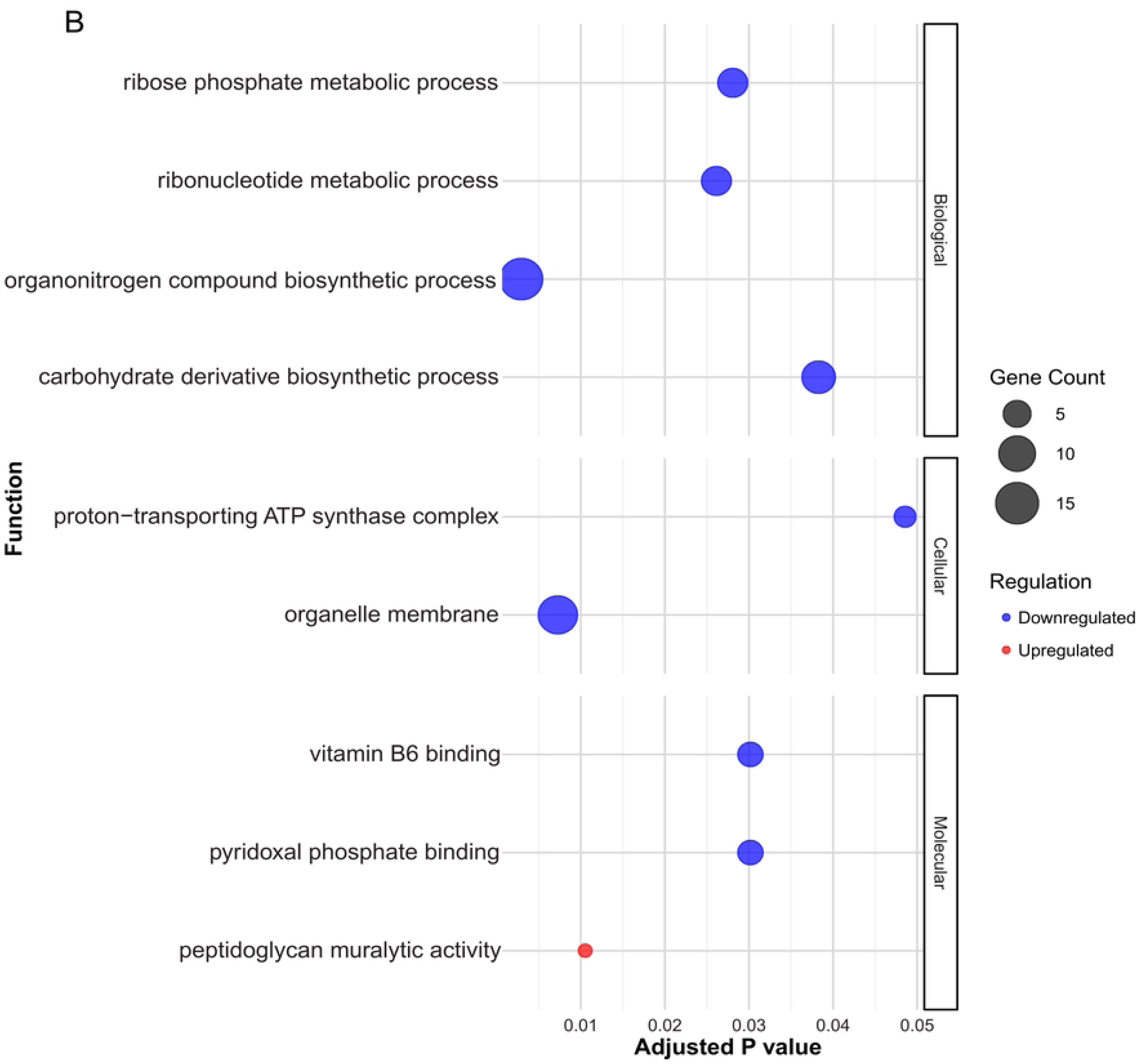
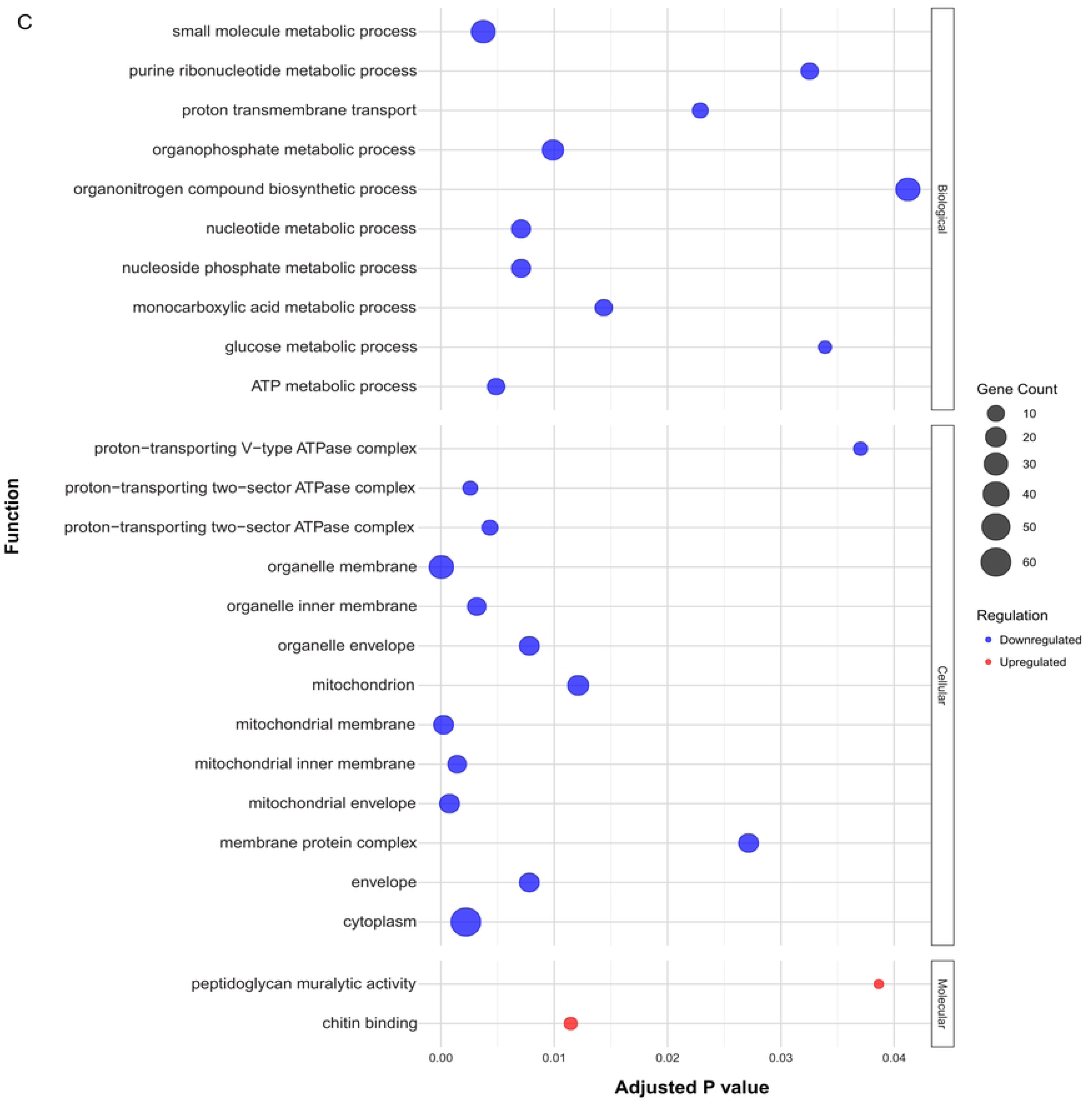
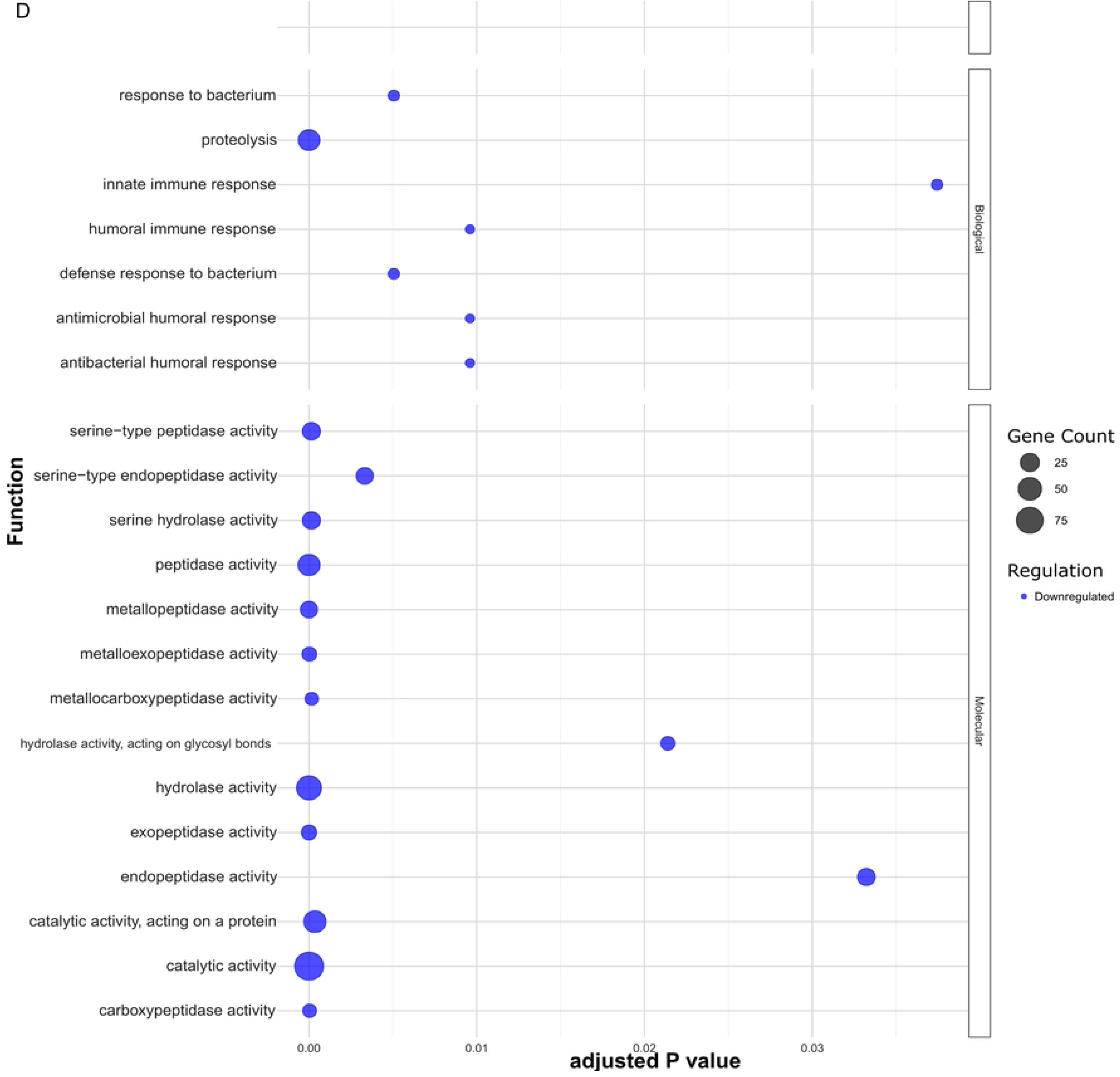
Beta diversity estimates of the gut microbiota data using unweighted unifrac (A,C and E), and non-metric multi-dimensional scaling (B,D and F) in *Microsporidia MB* positive mosquito guts (A and B), -negative pooled sibling mosquitoes (C and D) and -negative pooled non-sibling mosquitoes (E and F), across various time points 7 day old non-blood fed (yellow), 24 (green) and 48 hours (pink) post blood meal.

## Discussion

We demonstrate that *Microsporidia MB*-positive, non-blood-fed mosquitoes exhibited downregulation of immunity-related genes. Interestingly, immunity-related factors were upregulated following blood feeding, as shown by transcriptional profiling at 24- and 48-hours post-blood meal in both the gut and fat body tissues. Later, immune gene expression declined between 48- and 72-hours post-blood meal in both the gut and fat body. Strikingly, immune gene upregulation coincided with reduced expression of metabolic genes in *Microsporidia MB* positive mosquitoes. Notably, the downregulation of immunity-related genes in non-blood-fed *Microsporidia MB*-positive mosquitoes may shape the gut microbiota, promoting the presence of a denser and compositionally distinct community at the onset of blood feeding (when *P. falciparum* infection would occur). In addition, changes in immune-related gene upregulation at 24 and 48 hours may contrastingly alter the midgut microbiome after blood feeding. Altogether, this inconsistency with what could be considered a ‘normal’ *Anopheles* midgut could create a less permissive environment for *P. falciparum*.

We observed that gene expression and microbiota member composition in *Microsporidia MB*- positive and -negative pooled sibling mosquitoes clustered more closely, relative to the - negative pooled non-sibling mosquitoes. These findings suggest two possibilities: (1) genetic and/or environmental differences between *Microsporidia MB-*positive and -negative pooled non-sibling mosquitoes (rather than *Microsporidia MB* presence) are the paramount drivers of differences in gene expression and microbiota; or (2) *Microsporidia MB* induces transcriptional changes that persist even after the infection is cleared in pooled siblings. Since it is not fully possible to determine the importance of 1) vs. 2), we investigated and evaluated changes in gene expression in *Microsporidia MB*-positive mosquitoes relative to both *Microsporidia MB*- negative pooled sibling and pooled non-sibling mosquitoes.

### Transcriptional changes in the guts of non-blood-fed mosquitoes correlate with shifts in microbiota composition in *Microsporidia MB-*positive mosquitoes

It was evident that guts from *Microsporidia MB*-positive mosquitoes, compared to both *Microsporidia MB*-negative control groups, exhibited upregulated genes involved in digestion (*chymotrypsin 2-like*, *serine protease SP24D-like*) (18), juvenile hormone biosynthesis (*farnesoic acid O-methyltransferase*) (19), and detoxification (*glutathione S-transferase 1-1- like*) (20). Notably, *farnesol dehydrogenase* (21), another enzyme in the juvenile hormone biosynthesis pathway, displayed opposite expression trends: it was upregulated when comparing *Microsporidia MB*-positive mosquitoes to -negative pooled non-sibling mosquitoes but downregulated when compared to -negative pooled sibling mosquitoes. This suggests a potentially complex link between juvenile hormone and *Microsporidia MB*. For example, it is possible that *farnesol dehydrogenase* is not only linked to the presence of *Microsporidia MB* but also its clearing in the negative pooled sibling mosquitoes.

We also observed a downregulation of early *trypsins (trypsins 4, 6, and 7*) and concurrent upregulation of some *serine proteases*, which may play roles in both immunity and digestion. Previous studies have shown that juvenile hormone upregulation suppresses immune responses in mosquitoes (22), which is consistent with the observed downregulation of early *trypsins*, potentially involved in initiating the melanization cascade (23,24). These transcriptional changes may also selectively reshape the gut proteolytic environment, with potential consequences for microbiota dynamics (22,25,26). Specifically, digestive and detoxification enzymes may degrade or neutralize proteins and signaling molecules critical for bacterial colonization (25,26). Moreover, immune suppression linked to elevated juvenile hormone biosynthesis has been associated with increased microbial proliferation, enabling greater bacterial growth (22). Consistent with these patterns, *Microsporidia MB*-positive mosquitoes displayed distinct gut microbiota profiles and significantly elevated bacterial densities compared to both pooled sibling and pooled non-sibling controls. Together, these findings support the hypothesis that the upregulation of digestive, detoxification, and juvenile hormone biosynthesis pathways contributes to the modulation of gut microbiota in *Microsporidia MB*- infected mosquitoes.

Additionally*, Microsporidia MB* infection also influenced microbial abundances. Compared to both control groups, *Rhodococcus, Elizabethkingia, Chryseobacterium, Brevundimonas,* and *Mesorhizobium* were more abundant in *Microsporidia MB*-positive mosquitoes, while *Serratia* and *Pseudomonas*, which have anti-*Plasmodium* properties, were reduced. While several of these microbes play roles in digestion, oviposition attraction, and immune activation, *Rhodococcus* was particularly intriguing (27–32). In *Rhodnius prolixus*, *Rhodococcus rhodnii* serves as an endosymbiont, providing the host with vitamin B (28). In our dataset, although *Rhodococcus* was not likely endosymbiotic, its increased abundance in *Microsporidia MB*- positive mosquitoes suggests its potential role in vitamin B metabolism. Supporting this hypothesis, fat bodies of *Microsporidia MB*-positive mosquitoes exhibited increased metabolic activity compared to *Microsporidia MB-*negative pooled non-siblings, coinciding with the upregulation of vitamin B6 homeostasis pathways. This metabolic shift may be driven by an increased abundance of *Rhodococcus*, supplying cofactors for fat body metabolism—the central metabolic hub in insects.

Finally, *cysteine sulfinic acid decarboxylase*, consistently upregulated in the fat bodies of *Microsporidia MB*-positive mosquitoes compared to both *-*negative controls, may also play a role in enhanced metabolic responses in -positive mosquitoes. This enzyme contributes to taurine biosynthesis, which is essential for redox balance, osmoregulation, and metabolic fitness (33). These findings suggest that *Microsporidia MB* infection modulates mosquito physiology through a complex interplay of immune suppression, metabolism, and gut microbiota restructuring in non-blood fed mosquitoes.

### *Microsporidia MB* infection activates the immune system 24 hours post blood meal

During *Plasmodium* infection in mosquitoes, microgametes penetrate macrogametes in the midgut, forming zygotes. These zygotes develop into elongated, motile ookinetes within 22– 24 hours post-blood meal, establishing the infection (34). Ookinete invasion triggers a strong nitration response in the midgut and basal lamina, leading to hemocyte activation upon encountering the nitrated basal lamina. This immune priming induces the release of the hemocyte differentiation factor (HDF) into the hemolymph, increasing circulating hemocytes (35).

To investigate the potential impact of *Microsporidia MB* infection at this critical time point, we analyzed gene expression and gut microbiota shifts 24 hours post-blood meal. Our data indicates that *Microsporidia MB*-positive mosquitoes experienced reduced cellular stress (36), evidenced by the downregulation of *glycine-rich cell wall structural protein* and *cell wall integrity and stress response component 4-like* compared to -negative pooled sibling mosquitoes. Blood feeding is typically associated with cellular stress due to rapid temperature and pH shifts, gut distension, heme toxicity, and gut redox stress (37,38). Interestingly, *glutaredoxin C4* and *peroxiredoxin 6-like*, both of which function as antioxidants during oxidative stress (20,39), were upregulated in *Microsporidia MB*-positive mosquitoes compared to -negative pooled non-sibling mosquitoes, supporting the hypothesis that *Microsporidia MB* potentially alleviated cellular and oxidative stress after blood feeding.

Additionally, immune-related genes were upregulated, while metabolism-related genes were downregulated in *Microsporidia MB*-positive mosquitoes compared to negative pooled non-sibling mosquitoes. Immune-related genes included *peptidoglycan recognition protein SC* which is a pattern recognition receptor*, lysozyme C-1*, and *lipopolysaccharide tumor necrosis factor* (*LITAF*), as well as pathways such as the *peptidoglycan muralytic pathway* (40–42). Metabolism-related genes, such as *molybdenum cofactor sulfurase, serine palmitoyltransferase*, and *alanine glyoxylate aminotransferase*, were downregulated, along with pathways related to ribonucleotide metabolism, organonitrogen biosynthesis, carbohydrate derivatives, and Vitamin B6 binding.

While metabolism was generally downregulated in the gut of *Microsporidia MB*-positive compared to -negative pooled non-sibling mosquitoes, some processes such as peptide metabolic, cellular amide metabolic process, macromolecule metabolic process, were upregulated in the fat body of *Microsporidia MB*-positive compared to -negative pooled non-sibling mosquitoes. This may reflect the role of the fat body in vitellogenesis and energy provisioning to the ovary, where *Microsporidia MB* proliferates (43,44). The downregulation of pathways involved in cellular transport and mitochondrial process such as ATP metabolic, mitochondrion process, proton transport in the fat bodies of *Microsporidia MB*-positive compared to -negative pooled non-sibling mosquitoes, likely reflects a resource allocation shift towards immunity-related functions, which have high energy demands (45). Interestingly, oxidative stress associated genes such as *glutathione S-transferase* were downregulated in the fat body of *Microsporidia MB*-positive compared to -negative pooled non-sibling mosquitoes (20), further supporting the hypothesis that *Microsporidia MB*-positive mosquitoes maintain tight oxidative stress control, potentially minimizing excessive immune activation through processes such as melanization.

The upregulation of *LITAF*, which plays a role in hemocyte activation during *Plasmodium* infection in the fat body of *Microsporidia MB*-positive compared to -negative pooled non-sibling mosquitoes, suggests a systemic immune response extending beyond the gut. Activation of hemocyte differentiation likely enhances cellular and humoral immune responses, potentially contributing to *Plasmodium* transmission blocking (42,46,47). While cellular immunity appears central to this phenotype, the upregulation of *urease accessory protein* in *Microsporidia MB*-positive mosquitoes compared to both pooled sibling and pooled non-sibling mosquitoes suggests that gut environment modulation may also be involved. Since *urease* hydrolyzes urea into ammonia and carbon dioxide, its upregulation could alter gut pH, creating an inhospitable environment for *Plasmodium* parasites, as has been shown with carbonic anhydrase which was shown to modulate the gut pH of mosquitoes (48,49).

Furthermore, gut microbiota changes reinforce this hypothesis. The upregulation of the peptidoglycan muralytic pathway, which degrades Gram-positive bacteria (37), in *Microsporidia MB*-positive mosquitoes compared to -negative pooled non-sibling controls, suggests a shift favoring Gram-negative bacteria, which are known to enhance mosquito immunity against *Plasmodium*. *Serratia* and *Pseudomonas*, both of which have been linked to immune activation in response to *Plasmodium* infection (27,29,30), proliferated post-blood meal in *Microsporidia MB*-positive mosquitoes compared to both pooled sibling and pooled non-sibling controls. Since these bacteria possess urease activity, they may contribute to nitrogen balance alterations in the gut, enhancing the *Microsporidia MB* transmission-blocking effect (50,51).

Conversely, other microbes such as *Rhodococcus*, *Elizabethkingia*, *Chryseobacterium*, *Brevundimonas*, and *Mesorhizobium* declined significantly in *Microsporidia MB*-positive mosquitoes compared to both pooled sibling and pooled non-sibling controls. This decline suggests that these microbes may not be essential for mosquito physiology or *Plasmodium* transmission at this time point (28,31,32). Notably, the marked reduction in *Rhodococcus* could explain the downregulation of Vitamin B6 binding pathways, as this bacterium is associated with Vitamin B6 metabolism.

These findings collectively highlight how *Microsporidia MB* infection modulates immune responses, gut microbiota, and cellular stress, ultimately impairing *Plasmodium* transmission in mosquitoes.

### *Microsporidia MB* infected mosquitoes were marked by an activated immunity and slowed metabolism even at 48 hours post blood meal

After the ookinetes transverse the midgut epithelial barrier, they settle at the basal lamina of the gut where they develop into oocysts usually around 48 hours after a blood meal (34). We examined the effect of mosquito infection with *Microsporidia MB* at this time point and observed that there were no differences in gene expression profiles in the guts of *Microsporidia MB*-positive and -negative pooled sibling mosquitoes. Intriguingly, *cecropin B*, an antimicrobial peptide with anti-*Plasmodium* properties in mosquitoes (52), was downregulated in the fat body of *Microsporidia MB*-positive mosquitoes compared to -negative pooled sibling mosquitoes. While the magnitude of cecropin downregulation may not be in high orders of magnitude and have major consequences for *Plasmodium* refractoriness, the findings here suggest that *Microsporidia MB* potentially exhibits its primary effects on *Plasmodium* transmission blocking early after blood feeding among mosquitoes from *Microsporidia MB*-positive G_0_ females, and by 48 hours the effects have normalized, meaning initial immune activation and metabolic responses have returned to baseline levels

Interestingly, a comparison between *Microsporidia MB*-positive and -negative pooled non-sibling mosquitoes showed that *Microsporidia MB*-positive mosquitoes were still associated with an activated immune system, since the *peptidoglycan recognition SC2-like* and *lysozyme c-1* (40,41) were still upregulated at this time point. While immune system activation was evident, other innate immunity-related genes were generally downregulated, including *leucine rich melanocyte differentiation associated protein-like* and *galectin 4 like*, which is a pattern recognition molecule (53). Downregulation of some immune genes suggests a shift in immune focus at this time point in *Microsporidia MB*-positive mosquitoes compared to the negative pooled non-sibling mosquitoes.

Remarkably, unlike the downregulation of immunity when *Microsporidia MB*-positive mosquito fat bodies were compared to those from -negative pooled sibling mosquitoes, we observed an upregulation of *lysozyme c-1* in the fat body when *Microsporidia MB*-positive mosquitoes were compared to negative pooled non-sibling mosquitoes. The expression of this gene in the guts and fat body when *Microsporidia MB*-positive and -negative pooled non-sibling mosquitoes were compared suggests its systemic expression and immune system upregulation in *Microsporidia MB*-positive mosquitoes. This indicates a stronger innate immune response from *Microsporidia MB* infection compared to those without prior exposure. Strikingly, an upregulated immune system in our dataset was on the other hand associated with a downregulation in metabolic processes in both the fat body and the gut, further supporting previous evidence that immunity and metabolism exist in a compensatory dynamic (45).

Whereas *Serratia* and *Pseudomonas* generally recorded proliferation at the 24-hour time point, at the 48-hour time point they were marked by a drop in relative abundances to levels comparable to those in non-blood fed mosquitoes, while *Rhodococcus* and *Elizabethkingia* had their relative abundances rise again at this time point. The general reinstatement in these bacterial abundances to levels like those in non-blood fed mosquitoes support the gut transcriptomic evidence that this time point may be an inflection point to a slowed activation of immunity and an increased activation of metabolic processes in *Microsporidia MB*-positive compared to the -negative pooled non-sibling mosquitoes.

Evidence from this time point therefore suggests that offspring from G_0_ *Microsporidia MB*-positive females may have undergone immune priming leading to less aggressive responses compared to when comparisons are made between *Microsporidia MB*-positive and -negative pooled non-sibling mosquitoes.

### *Microsporidia MB* infected mosquitoes associated with downregulated immune system and upregulated metabolism genes 72 hours post blood meal

This time point coincides with the progressive development of the *Plasmodium* oocysts before their maturation and release as sporozoites after 7-14 days post a blood meal (34). This time point was associated with upregulated metabolic genes, including *acetyl-CoA carboxylase*, *homocysteine S-methyl transferase*, *molybdenum cofactor sulfurase*, and *fatty acid synthase* (54–56), when guts from *Microsporidia MB*-positive mosquitoes were compared to -negative pooled sibling mosquitoes. When *Microsporidia MB*-positive and -negative pooled non-sibling mosquito guts were compared, we observed downregulated immunity genes in *Microsporidia MB*-positive mosquitoes, including *defensin*, *cecropin A*, *cecropin B*, *cecropin C*, *lipopolysaccharide induced tumor necrosis factor*, *peptidoglycan recognition protein* and *lysozyme c like* (46,47,57,58). Upregulated metabolism in *Microsporidia MB*-positive mosquitoes may suggest a faster metabolic recovery in these mosquitoes compared to *Microsporidia MB-*negative pooled sibling mosquitoes at this time point. Downregulated immune system activation at this time point when comparing *Microsporidia MB*-positive and -negative non-sibling mosquitoes may suggest a wane in the immune system activation from this time point onwards at the transcriptomics level.

In the fat body, *ornithine decarboxylase* (*ODC*) and *galectin* were downregulated when *Microsporidia MB*-positive mosquitoes were compared to -negative pooled sibling mosquitoes. *ODC* has an associated role in polyamine biosynthesis (59). Its downregulation alongside *galectin* which is a pattern recognition receptor (53), support a shift from immune activation and metabolism of polyamines elevated by blood feeding in *Microsporidia MB*-positive mosquitoes. A comparison of the fat bodies from *Microsporidia MB*-positive and -negative pooled non-sibling mosquitoes was also associated with an upregulation of metabolism related pathways and genes, including ATP synthase subunit, purine nucleoside phosphorylase, bifunctional purine biosynthetic protein and phosphoenolpyruvate carboxykinase.

The general observation of a waning immune system activation at this time point in *Microsporidia MB*-positive mosquitoes suggests immune system stabilization which is associated with diversion of resources to maintaining cellular processes and functions. The striking downregulation of immune system activation even when *Microsporidia MB*-positive mosquitoes are compared to *Microsporidia MB*-negative pooled sibling mosquitoes supports the idea that *Microsporidia MB*-positive mosquitoes have a primed immune system that may be entering a state of less activation due to a maintained balanced state to avoid overreaction at this time point. These findings further support the conclusion that *Plasmodium* transmission blocking using the endosymbiont *Microsporidia MB* happens earlier, after blood feeding.

## Conclusion

Our results suggest that *Microsporidia MB*-infected mosquitoes have a slightly lowered ‘resting’ midgut immune activity when non-blood fed. Once the mosquito takes a blood meal, there seem to be a potential ‘over-compensation’ with elevated gene expression in immune-related genes relative to *Microsporidia MB* uninfected mosquitoes. The changes in immune dynamics over the period of potential infection with *Plasmodium* could promote refractoriness if the mosquito midgut environment becomes less conducive at critical stages in the *Plasmodium* infection cycle (Figure 8). *Microsporidia MB* may also modulate the midgut physiology (e.g pH) making the midgut environment unconducive for *Plasmodium* development. Lastly, *Microsporidia MB* may also impair the transmission of *Plasmodium* by altering the gut microbiota composition (either directly or via physiological or immunological changes) in infected mosquitoes, promoting microbes with anti-*Plasmodium* properties.

**Figure 8:**
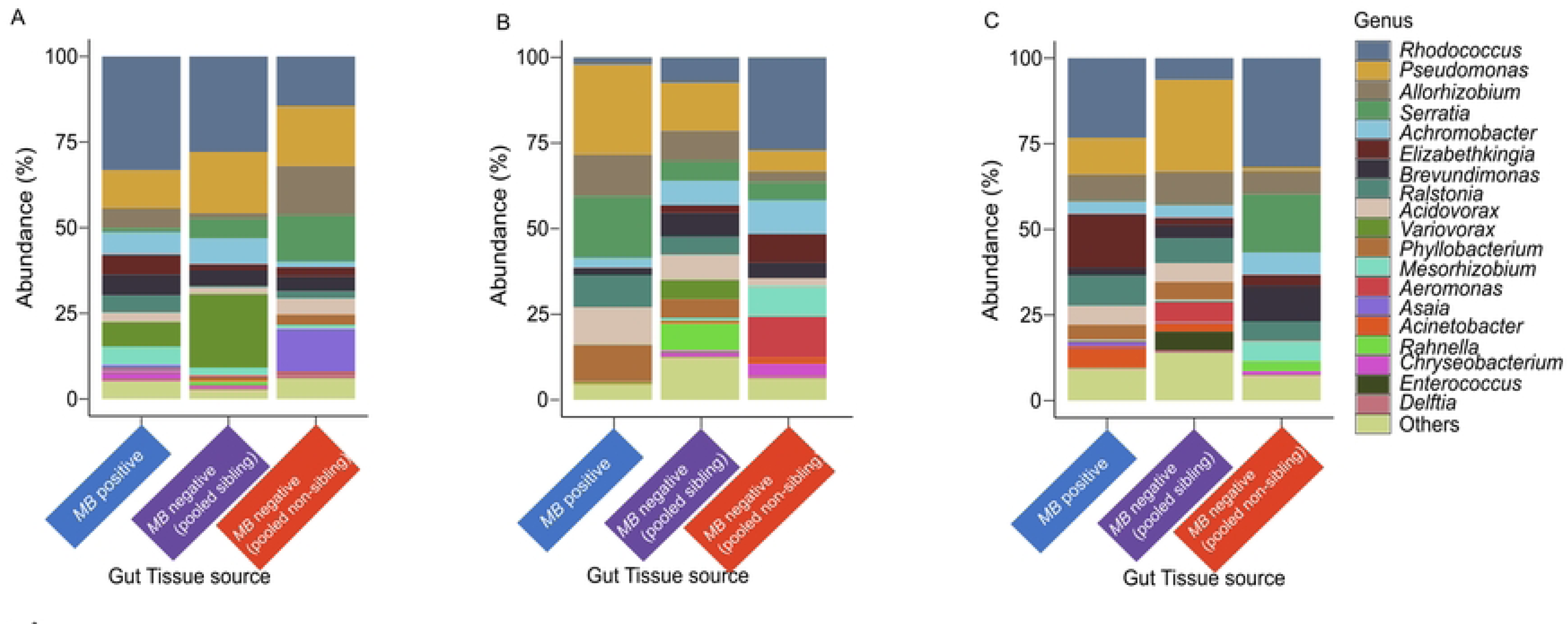
Proposed model for *Plasmodium falciparum* transmission blocking using the endosymbiont *Microsporidia MB***. 1. Microbiota**: Non-blood fed mosquitoes demonstrated a distinct gut microbiota structure and a downregulation of the immune system favoring further proliferation of gut microbes, which could be creating a favorable environment that faces the arrival of *Plasmodium* with blood feeding. In addition, it was evident that blood feeding potentially favored the modulation of the pH in the gut and the proliferation of microbes with anti-*Plasmodium* properties 24 hours post blood meal, including *Serratia* and *Pseudomonas* which are also associated with activating the host immune system. **2. Hemocyte differentiation**: the immune system in *Microsporidia MB* infected mosquitoes was activated with blood feeding, with evidence of factors with roles in hemocytes differentiation being activated. This indicates the potential role of the cellular immune responses in *Plasmodium* transmission blocking in mosquitoes infected with *Microsporidia MB*.

## Methods

### Ethical approval

The Kenya Medical Research Institute Scientific Review Unit and the University of Cape Town Human Research Ethics Committee granted ethical approval to support this study under protocol numbers **KEMRI/RES/7/3/1** and **HREC REF: 663/2023**, respectively.

### Description of the study samples

Gravid female *An. arabiensis* mosquitoes were manually aspirated from inside walls of houses neighboring the Ahero irrigation scheme (-34.9190W, -0.1661N) in Kenya. The wild caught G_0_ female mosquitoes were induced to oviposit inside perforated 1.5 ml microcentrifuge tubes containing 50 µl of distilled water and a soaked Whatman paper (1 cm × 1 cm in size) (7). Upon laying eggs, these females were then screened for the presence of *Microsporidia MB* by targeting the *Microsporidia MB* 18S gene (Microsporidia MB and Blood Meal Cytochrome b Screening). using polymerase chain reaction (PCR). Eggs from each confirmed *Microsporidia MB*-positive and -negative G_0_ wild-caught female *An. arabiensis* mosquitoes were individually transferred to water trays for larval development under standardized conditions (27 ± 2.5°C, humidity 60–80%), with negative females serving as experimental controls (7). Eggs from both *Microsporidia MB*-positive and -negative G_0_ *An. arabiensis* females were dispensed into separate trays for rearing at the larval stage and pupae collected in pools after pupation and transferred into separate cages (those from positive and negative G_0_ females in different cages) for adult emergence and adult *An. arabiensis* first generation (G_1_) rearing. G_1_ adult rearing cages were divided into two experimental conditions as follows: (i) those positive for *Microsporidia MB*, and (ii) those negative for *Microsporidia MB* as determined by screening for the *Microsporidia MB* 18S gene of the G_0_ *An. arabiensis* females (Microsporidia MB and Blood Meal Cytochrome b Screening). Negative mosquitoes from positive G_0_ females are termed negative pooled sibling mosquitoes while negative mosquitoes from negative G_0_ females are termed negative pooled non sibling mosquitoes as pupae from various trays were pooled for adult emergence.

Seven-day old non-blood-fed adult mosquitoes and 4-, 5- and 6-day-old blood-fed (to represent 24, 48, and 72 hours post blood meal, respectively) adults fed on blood from mice at three days post-emergence were dissected from the positive and control groups to obtain guts, ovaries and fat bodies. Screening for blood feeding was done by targeting the Cytochrome b (*Cyt b*) gene from the midguts of blood-fed mosquitoes (Microsporidia MB and Blood Meal Cytochrome b Screening).

### Transcriptomics profiling

#### *RNA* and *DNA* extraction

Trizol reagent (Invitrogen) was used to extract *RNA* and *DNA* from the different mosquito samples spanning the two experimental conditions in the study (G_1_ from Microsporidia MB-positive and negative mothers). Briefly, the *RNA* phase was isolated using chlorofoam, after which *RNA* was precipitated using isopropanol. *DNA,* on the other hand, was extracted from the organic phase using the back extraction buffer (BEB) [4M Guanidine thiocyanate, 50mM sodium citrate, 1M Tris (free base)] and precipitated using isopropanol. Two 75% ethanol wash steps were done during *RNA* extraction while during *DNA* extraction two 70% ethanol wash steps were done, followed by resuspension of both nucleic materials in nuclease-free water. The purity of both *RNA* and *DNA* was determined using NanoDrop One/Oneᶜ Microvolume UV-Vis Spectrophotometer (ThermoFischer Scientific, Waltham, Massachusetts, United States), while quantification of both *RNA* and *DNA* was done using Qubit™ 4 Fluorometer (Invitrogen, Waltham, Massachusetts, United States) throughout the entire study. For each sample, 50 ng of the extracted *RNA* was prepared for sequencing using the Oxford Nanopore Technology (ONT) MinION device. The *DNA* from the corresponding samples was used for screening of *Microsporidia MB* using PCR, and for determining the relative ratios of *Microsporidia MB* and host DNA present in the ovary samples (Microsporidia MB and Blood Meal Cytochrome b Screening) Samples were classified as *Microsporidia MB*-positive throughout the study based on detection of infection in the ovaries.

### Complementary DNA (cDNA) synthesis and *RNA* sequencing

A total of 96 samples were used for *RNAseq,* where 24 replicates were considered per experimental condition. *RNA* samples that passed the quality assessment and possessed good concentrations were converted to cDNA by targeting the *polyA* tail of the *mRNA* using VN primers present in the PCR cDNA barcoding (SQK-PCB109) (Oxford Nanopore Technologies) kit. Maxima H Minus Reverse Transcriptase (ThermoFischer Scientific) and the Strand Switching Primer (PCR cDNA barcoding kit) were also used during the cDNA synthesis process in an incubation reaction at 42 °C for 90 min, followed by heat inactivation at 85 °C. Next, the barcodes in the PCR cDNA barcoding kit were used for amplification and barcoding of the samples. The LongAmp Taq Master Mix (New England BioLabs) was used for the amplification reaction with the PCR conditions set as follows: initial denaturation at 95 °C for 30 s, denaturation at 95 °C for 15 s, annealing at 62 °C for 15 s for 11-18 cycles, extension at 65 °C for 50 s for 11-18 secs and final extension 65 °C for 6 min. The PCR product was cleaned using the Agencourt Ampure XP beads (Beckman Coulter) to remove unused primers, partial amplicons, and enzymes, followed by adapter ligation. 37.5 µl Sequencing Buffer and 25.5 µl Loading Beads (PCR cDNA barcoding-SQL-PCB109) were then added to the *DNA* library, followed by loading of the library on the ONT MinION device for sequencing.

### Bioinformatics analysis and differential gene expression analysis

*RNA*seq data basecalling and demultiplexing were done live using the Guppy software (Version 6.5.7)(60) . A meta-transcriptomics approach was used in data analysis as follows: data quality was determined using PycoQC (Version 2.5.2) (61) followed by trimming of adapters using porechop (Version 0.2.4) (62). Residual ribosomal *RNA* was then removed using SORTMERNA (Version 4.3.4) (63), after which cluster gene isoform generation using IsONclust (Version 0.0.9) was done (64). The errors in the resultant reads were corrected using IsONcorrect (Version 0.0.3), and the error corrected *mRNA* was then aligned to the corresponding *An. arabiensis* transcriptome present in VectorBase (Release 67) (*Anopheles arabiensis* Dongola 2021, Species NCBI Taxon ID 7173) using minimap2 (Version 2.24) (65). Transcript quantification was done using Nanocount (Version 1.1.0.post6) (66), and the recovered transcripts annotated using VectorBase for identification. Next, differential expression analysis beginning with the normalization of the library was done in R (Version 4.4.1) using the edgeR package (Version 4.2.1) (67). Samples were included in downstream statistical analysis if they recorded comparable read counts with the rest of the samples in the comparison groups, and if they didn’t cluster too distinctly using principal component analysis, rendering their inconsideration as outliers. Based on this inclusion criteria, out of the 96 samples included in the transcriptomics analysis, four including sample_8FB, sample_34FB, sample_56FB and sample_92G were excluded from downstream analysis due to few read counts or consideration as outliers. On the other hand, genes were differentially expressed when the log fold change in the gene expression was >2, and the adjusted *P*-value was < 0.05. Gene enrichment analysis was also done in VectorBase to identify the gene ontologies of the differentially expressed genes.

### Gut microbiota profiling

#### DNA extraction

The various dissected tissues from the two different conditions (G_1_ from *Microsporidia MB*- positive and *-*negative mothers) and three time points (non-blood fed, 24 and 48 hours post blood meal) had their DNA extracted using the ammonium acetate protein precipitation method described in (7). DNA from the dissected tissues from *Microsporidia MB*-infected mothers were then screened to determine the *Microsporidia MB* infection status of the mosquitoes. Samples were also classified as *Microsporidia MB*-positive based on detection of infection in the ovaries.

#### *Microsporidia MB* and Blood Meal Cytochrome b Screening

*Microsporidia MB*-specific primers (MB18SF: CGCCGGCCGTGAAAAATTTA and MB18SR: CCTTGGACGTGGGAGCTATC) that target the *Microsporidia MB 18S* rRNA gene region were used in screening for *Microsporidia MB* positive ovary samples. Blood meal analysis, on the other hand, was done by screening the *Cyt b* mtDNA from the midguts of blood-fed mosquitoes using *Cytb* specific primers (FWD: GAGGMCAAATATCATTCTGAGG and REV: TAGGGCVAGGACTCCTCCTAGT). Conventional PCR reaction volume of 10 µl constituted of 2 µl HOTFirepol^®^ Blend Master mix Ready-To-Load (Solis Biodyne, Estonia, mix composition: 7.5 mM Magnesium chloride, 2 mM of each dNTPs, HOT FIREPol^®^ DNA polymerase), 0.5 µl of 5 pmol µl^−1^ of both forward and reverse primers, 2 µl of the template and 5 µl nuclease-free PCR water were prepared for *Microsporidia MB* and *Cytb* screening. The PCR cycling conditions were set as follows: initial denaturation at 95 °C for 15 min, further denaturation at 95 °C for 30 s, followed by annealing at 62 °C in the case of *Microsporidia MB* and at 58 °C in the case of *Cytb* for 30 s and extension at 72 °C for a further 30 s, all done for 35 cycles. The final extension was done at 72 °C for 5 min.

To determine *Microsporidia MB* and *Cytb* infection densities, *Microsporidia MB* and *Cytb* were quantified by *qPCR* using MB18SF/ MB18SR and *Cytb*F/*Cytb*R primers, respectively, with normalization against the *Anopheles* ribosomal *S7* host gene (primers, S7F: TCCTGGAGCTGGAGATGAAC and S7R: GACGGGTCTGTACCTTCTGG) using the same PCR conditions as those for the *Microsporidia MB* 18S region. 16S rRNA densities from the guts spanning the different experimental conditions were also determined by amplifying the V1-V2 region of the genes using (16S_27Fmod: TCG TCG GCA GCG TCA GAT GTG TAT AAG AGA CAG AGR GTT TGA TYM TGG CTC AG) and (16S_338R: GTC TCG TGG GCT CGG AGA TGT GTA TAA GAG ACA GTG CTG CCT CCC GTA GGA GT) primers (68), with the cycling conditions set as initial denaturation at 95 °C for 15 min, further denaturation at 95 °C for 30 s, followed by annealing at 63 °C for 30 s and extension at 72 °C for a further 30 s, all done for 35 cycles. The final extension was done at 72 °C for 5 min. Normalization for the bacterial densities in the guts of the mosquitoes was also done against the *Anopheles* ribosomal *S7* host gene. All qPCR assays in this study were carried out on the MIC qPCR cycler (Bio Molecular Systems, Australia).

### 16S rRNA gene amplification, sequencing and analysis

A total of 90 samples spanning the two experimental conditions (G_1_ from *Microsporidia MB*- positive and *-*negative G_0_ *An. arabiensis* females) in this experiment were shipped to Macrogen Europe BV (Meibergdreef, Amsterdam, Netherlands) for 16S rRNA amplification and sequencing (10 samples from 10 individual guts per treatment condition). The 16S rRNA region was amplified by targeting the V3-V4 region using the 314F- CCTACGGGNGGCWGCAG and 805R-GACTACHVGGGTATCTAATCC primers. After successful amplification, libraries for each sample were made, followed by sequencing on the Illumina MiSeq sequencing platform.

The 16S rRNA paired-end sequences spanning the V3-V4 region of bacteria were analyzed using the QIIME2 software (Version 2024.5) (69). Following read quality assessment and primer and adapter trimming using Cutadapt embedded in QIIME2, forward and reverse reads were merged, and chimeras were removed using the DADA2 pipeline in QIIME2. Taxonomic classification was then done against the Silva 138 database using a pre-trained naïve Bayes classifier. Subsequently, the resulting feature, taxonomy, and sequence tables from the QIIME2 pipeline were read into R for downstream statistical analysis. Amplicon sequence variants (ASVs) classified as Mitochondria, Chloroplast, Archaea, Cyanobacteria, Chloroflexi, Eukaryota and classified as unassigned reads at the kingdom level were filtered out from the analysis as a first step in R. Next, ASVs not classified up to the genus level and those with ambiguous classification had their sequences filtered out as a second step in R and re-classified using the NCBI blast 16S rRNA database (2020 release). After this classification, the results were merged to the silva classified results, and a Phyloseq object was created, followed by filtering ASVs with abundance below 5. Further downstream analyses to generate an abundance barplot of the most abundant bacteria in the different treatments, beta diversity estimates using unweighted unifrac and non-metric multidimensional scaling (NMDS), prevalence estimates, and PERMANOVA estimation were done using the R statistical software.

## Acknowledgement

The authors acknowledge Dr. Thomas Onchuru, Dr Rehemah Gwokyalya, Joseph Muthoni, and Fidel Gabriel Otieno for their incredible support with mosquito dissections and advice while performing all the experiments. We also thank Jeniffer Thiong’o, Peris Waweru and Mary Mwangi of the *icipe* Arthropod Rearing and Containment Unit for mosquito rearing assistance. We sincerely appreciate Faith Kyengo, Rose Marubu and Dr Thomas Onchuru for their administrative support and help with procurement of all reagents used in this study. The authors gratefully acknowledge the financial support for this research by the following organizations and agencies: Bill & Melinda Gates Foundation (INV0225840); the Swedish International Development Cooperation Agency (Sida); the Swiss Agency for Development and Cooperation (SDC); the Australian Centre for International Agricultural Research (ACIAR); the Government of Norway; the German Federal Ministry for Economic Cooperation and Development (BMZ); and the Government of the Republic of Kenya. The views expressed herein do not necessarily reflect the official opinion of the donors

## Supplementary information

**Figure S1:**
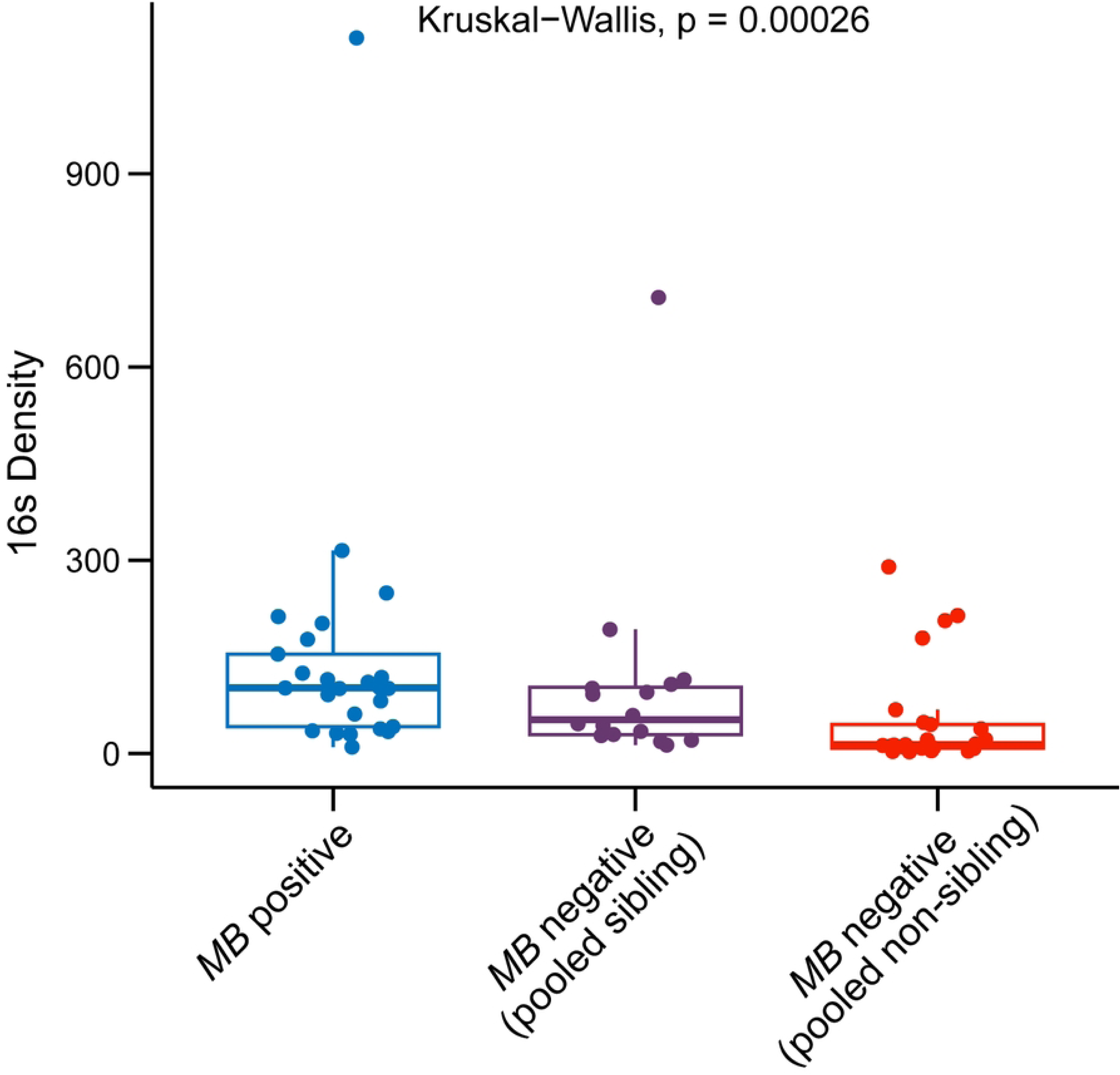
PCA to demonstrate the clustering patterns of genes from the fat bodies of *Microsporidia MB* positive compared to both negative pooled sibling and pooled non-sibling mosquitoes across different time points. (A) represents the clustering patterns in non-blood fed mosquitoes, (B) 24 hours post blood meal, (C) 48 hours post blood meal and (D) 72 hours post a blood meal. Blue represents *Microsporidia MB* positive mosquitoes, purple negative pooled sibling mosquitoes and red negative pooled non-sibling mosquitoes. Red represents upregulation while blue represents downregulation.

**Figure S2:**
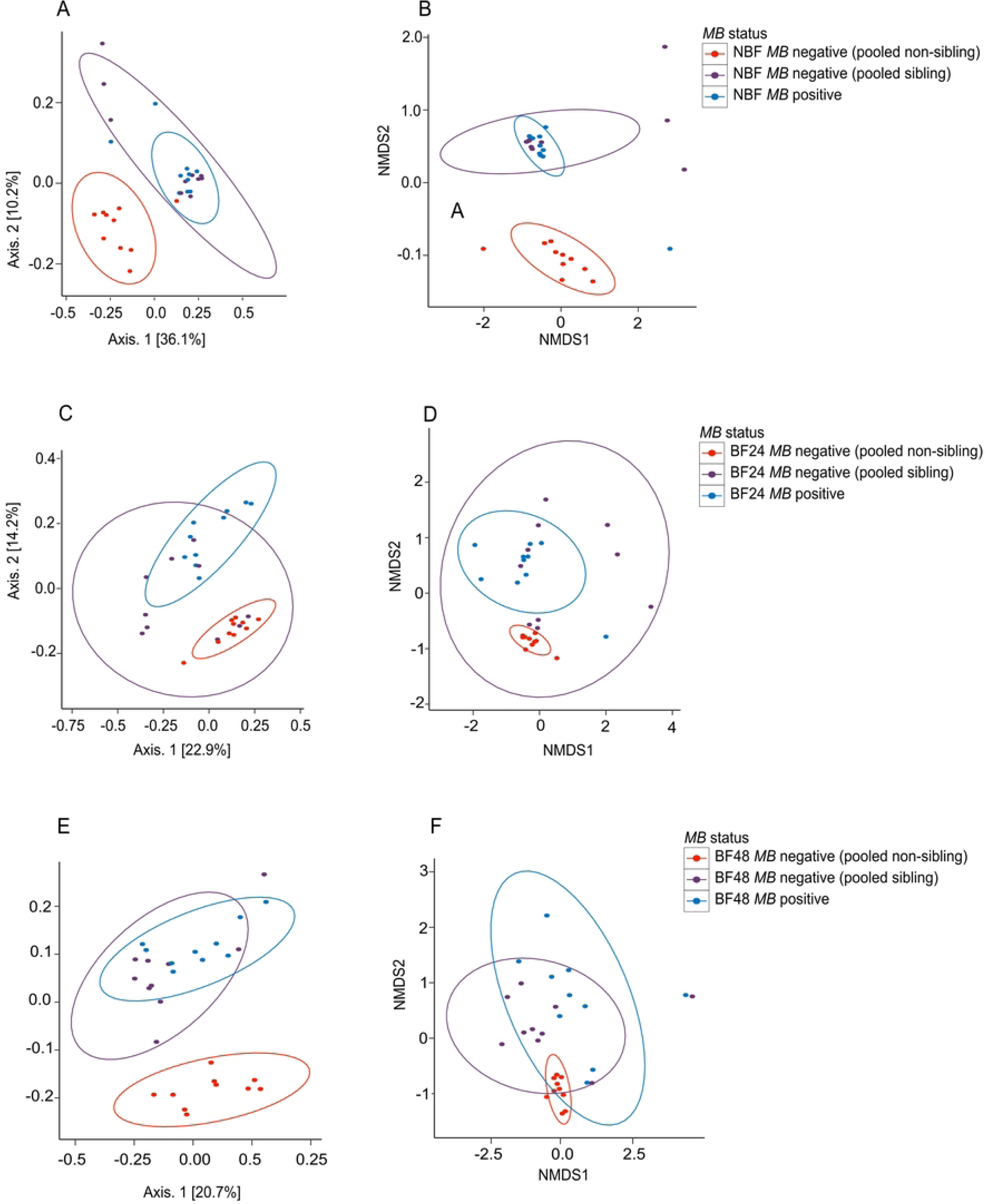
Volcano plots showing the differentially expressed genes in the fat bodies of *Microsporidia MB* positive mosquitoes compared to *Microsporidia MB* negative pooled non-sibling mosquitoes. A, B, C and D represent the differentially expressed genes in non-blood-fed mosquitoes, 24 hours post blood meal, 48 hours post blood meal and 72 hours post blood meal respectively. Red represents upregulation while blue represents downregulation.

**Figure S3:**
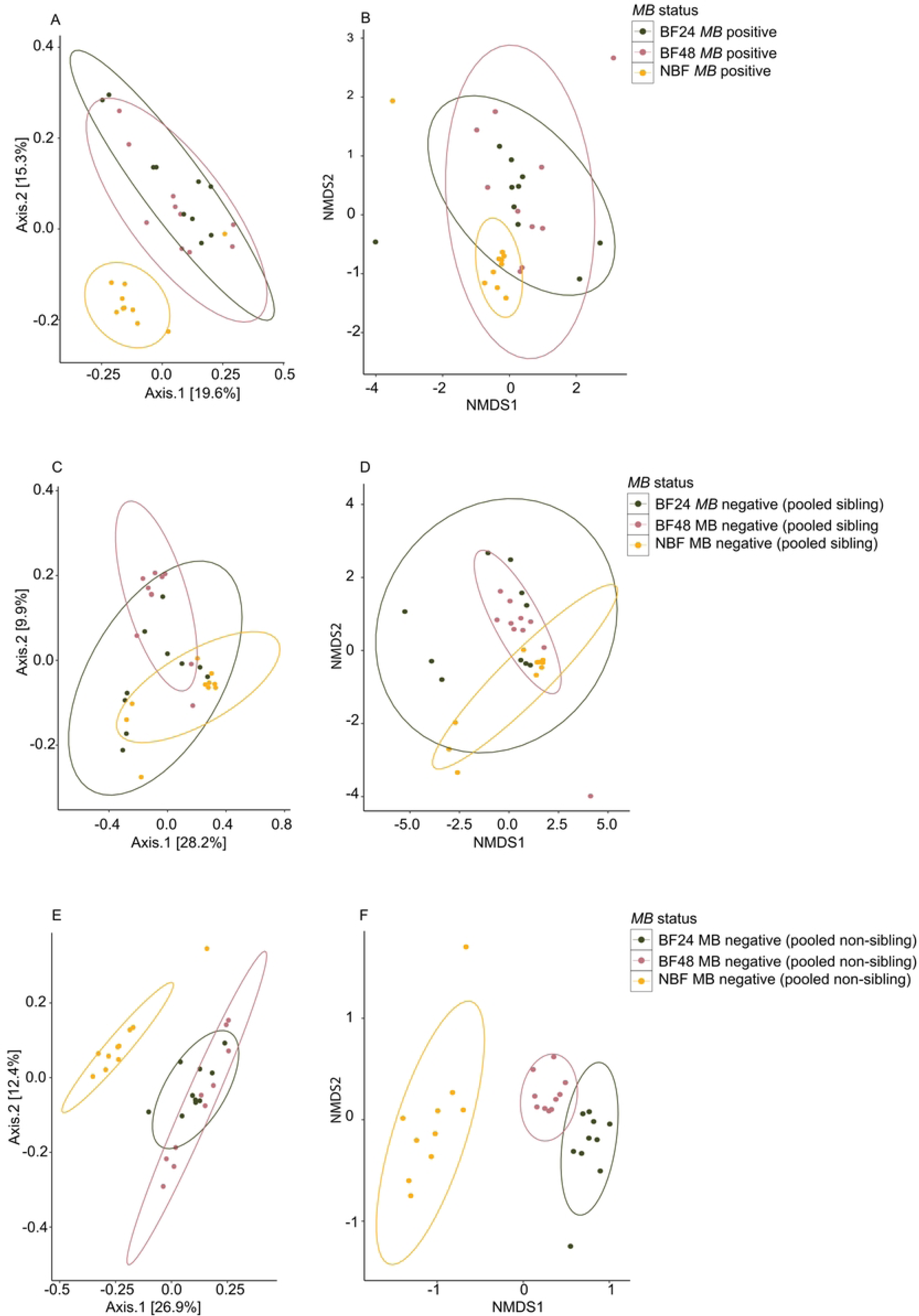
Gene ontology analysis of differentially expressed genes in the fat bodies of *Microsporidia MB* positive compared to negative pooled non-sibling mosquitoes. A, B, C and D represent the enriched pathways in non-blood-fed mosquitoes, 24 hours post blood meal, 48 hours post blood meal and 72 hours post blood meal respectively. Red represents upregulation while blue represents downregulation.

**Figure S4:**
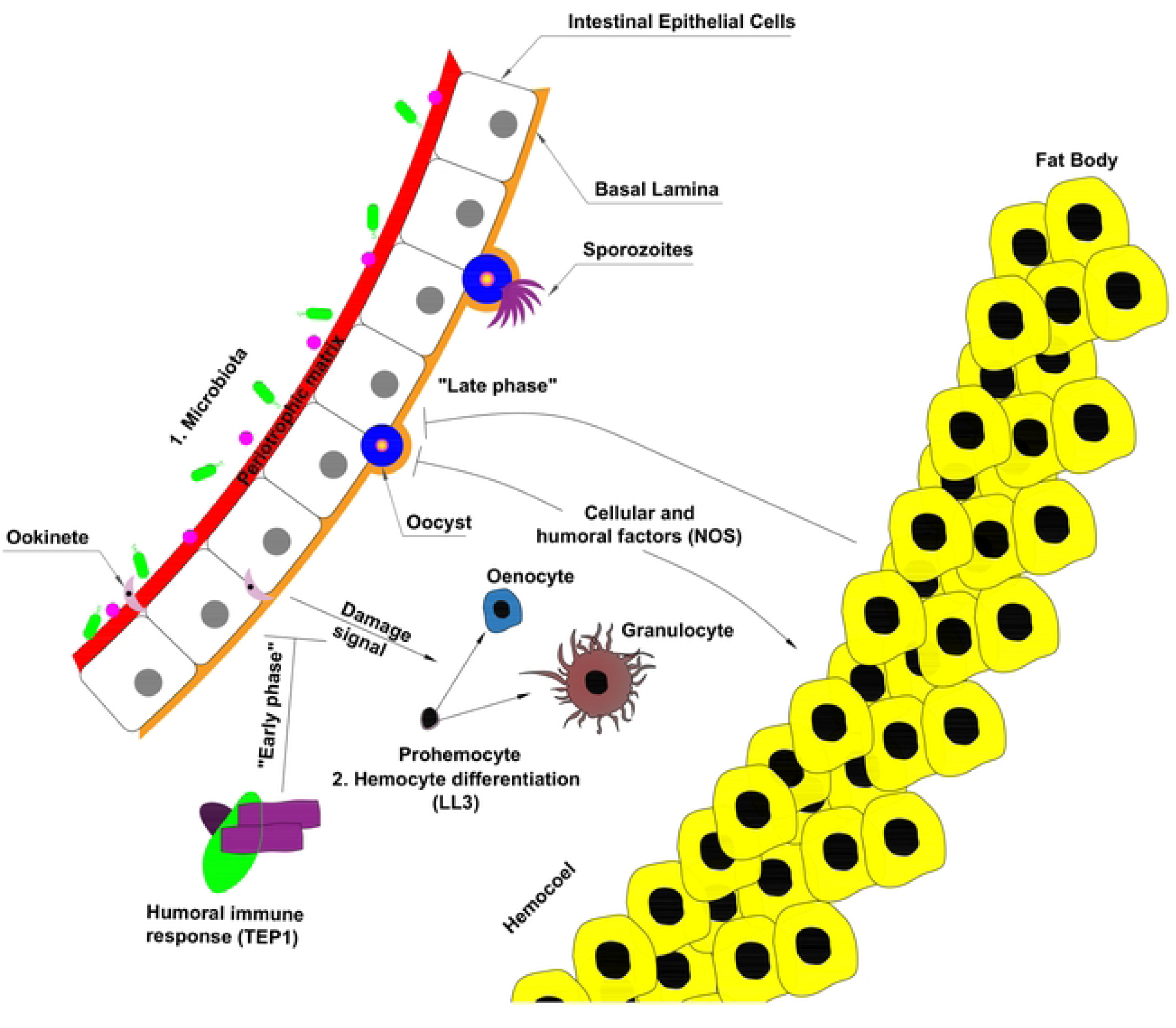
A heatmap showing the time series trends in the gene expression of some key genes in *Microsporidia MB* positive mosquito guts. Red represents upregulation while blue represents downregulation. Asterisks represent significant false discovery rates, with two asterisks indicating significance below 0.01 while one asterisk represents significance below 0.05

**Supplementary_table_S1**: Differentially expressed genes in non-blood fed *Microsporidia MB* positive *An. arabiensis* guts compared to -negative pooled sibling mosquitoes.

**Supplementary_table_S2**: Differentially expressed genes in *Microsporidia MB* positive *An. arabiensis* guts compared to -negative pooled sibling mosquitoes 24 hours post blood feeding

**Supplementary_table_S3**: Differentially expressed genes in *Microsporidia MB* positive *An. arabiensis* guts compared to -negative pooled sibling mosquitoes 48 hours post blood feeding

**Supplementary_table_S4**: Differentially expressed genes in *Microsporidia MB* positive *An. arabiensis* guts compared to -negative pooled sibling mosquitoes 72 hours post blood feeding

**Supplementary_table_S5**: Differentially expressed genes in non-blood fed *Microsporidia MB* positive *An. arabiensis* guts compared to -negative pooled non-sibling mosquitoes.

**Supplementary_table_S6**: Differentially expressed genes in *Microsporidia MB* positive *An. arabiensis* guts compared to -negative pooled non-sibling mosquitoes 24 hours post blood feeding

**Supplementary_table_S7**: Differentially expressed genes in *Microsporidia MB* positive *An. arabiensis* guts compared to -negative pooled non-sibling mosquitoes 48 hours post blood feeding

**Supplementary_table_S8**: Differentially expressed genes in *Microsporidia MB* positive *An. arabiensis* guts compared to -negative pooled non-sibling mosquitoes 72 hours post blood feeding

**Supplementary_table_S9**: Differentially expressed genes in non-blood fed *Microsporidia MB* positive *An. arabiensis* fat bodies compared to -negative pooled sibling mosquitoes.

**Supplementary_table_S10**: Differentially expressed genes in *Microsporidia MB* positive *An. arabiensis* fat bodies compared to -negative pooled sibling mosquitoes 24 hours post blood feeding

**Supplementary_table_S11**: Differentially expressed genes in *Microsporidia MB* positive *An. arabiensis* fat bodies compared to -negative pooled sibling mosquitoes 48 hours post blood feeding

**Supplementary_table_S12**: Differentially expressed genes in *Microsporidia MB* positive *An. arabiensis* fat bodies compared to -negative pooled sibling mosquitoes 72 hours post blood feeding

**Supplementary_table_S13**: Differentially expressed genes in non-blood fed *Microsporidia MB* positive *An. arabiensis* fat bodies compared to -negative pooled non-sibling mosquitoes.

**Supplementary_table_S14**: Differentially expressed genes in *Microsporidia MB* positive *An. arabiensis* fat bodies compared to -negative pooled non-sibling mosquitoes 24 hours post blood feeding

**Supplementary_table_S15**: Differentially expressed genes in *Microsporidia MB* positive *An. arabiensis* fat bodies compared to -negative pooled non-sibling mosquitoes 48 hours post blood feeding

**Supplementary_table_S16**: Differentially expressed genes in *Microsporidia MB* positive *An. arabiensis* fat bodies compared to -negative pooled non-sibling mosquitoes 72 hours post blood feeding

**Supplementary_table_S17**: Enriched Gene Ontology (GO) terms among differentially expressed genes in non-blood fed *Microsporidia MB* positive *Anopheles arabiensis* guts compared to -negative non-sibling mosquito guts

**Supplementary_table_S18**: Enriched Gene Ontology (GO) terms among differentially expressed genes in non-blood fed *Microsporidia MB* positive *Anopheles arabiensis* fat bodies compared to -negative non-sibling mosquito fat body

**Supplementary_table_S19**: Enriched Gene Ontology (GO) terms among differentially expressed genes 24 hours post blood meal in Microsporidia *MB* positive *Anopheles arabiensis* guts compared to -negative non-sibling mosquito guts

**Supplementary_table_S20**: Enriched Gene Ontology (GO) terms among differentially expressed genes 24 hours post blood meal in Microsporidia *MB* positive *Anopheles arabiensis* fat body compared to -negative non-sibling mosquito fat body

**Supplementary_table_S21**: Supplementary_table_S19: Enriched Gene Ontology (GO) terms among differentially expressed genes 48 hours post blood meal in Microsporidia *MB* positive *Anopheles arabiensis* guts compared to -negative non-sibling mosquito guts

**Supplementary_table_S22**: Enriched Gene Ontology (GO) terms among differentially expressed genes 48 hours post blood meal in Microsporidia *MB* positive *Anopheles arabiensis* fat body compared to -negative non-sibling mosquito fat body

**Supplementary_table_S23**: Supplementary_table_S19: Enriched Gene Ontology (GO) terms among differentially expressed genes 72 hours post blood meal in Microsporidia *MB* positive *Anopheles arabiensis* guts compared to -negative non-sibling mosquito guts

**Supplementary_table_S24**: Enriched Gene Ontology (GO) terms among differentially expressed genes 72 hours post blood meal in Microsporidia *MB* positive *Anopheles arabiensis* fat body compared to -negative non-sibling mosquito fat body

**Supplementary_table_S25**: Relative abundances of the most abundant bacteria in the guts of *An. arabiensis* mosquitoes across treatment groups

**Supplementary_table_S26**: A tabulation of the prevalence of bacteria in the guts of *An. arabiensis* mosquitoes included in the study

**Supplementary_table_S27**: Pairwise PERMANOVA results comparing microbial community composition across treatment groups.

## Notes

### Competing Interest Statement

The authors have declared no competing interest.

